# Interaction of a photochromic UV sensor protein Rc-PYP with PYP-binding protein

**DOI:** 10.1101/2021.06.01.446672

**Authors:** Suhyang Kim, Yusuke Nakasone, Akira Takakado, Yoichi Yamazaki, Hironari Kamikubo, Masahide Terazima

**Affiliations:** Department of Chemistry, Graduate School of Science, Kyoto University, Kyoto 606-8502, Japan; Division of Materials Science, Graduate School of Science and Technology, Nara Institute of Science and Technology, Takayama 8916-5, Ikoma city, Nara, 630-0192 Japan

## Abstract

Photoactive yellow protein (PYP) from *Halorhodospira halophila* is one of typical light sensor proteins. Although its photoreaction has been extensively studied, no downstream partner protein has been identified to date. In this study, the intermolecular interaction dynamics observed between PYP from *Rhodobacter capsulatus* (Rc-PYP) and a possible downstream protein, PYP-binding protein (PBP), were studied. It was found that UV light-induced a long-lived product (pUV*), which interacts with PBP to form a stable hetero-hexamer (Complex-II). The reaction scheme for this interaction was revealed using transient absorption and transient grating methods. Time-resolved diffusion detection showed that a hetero-trimer (Complex-I) is formed transiently, which produced Complex-II via a second-order reaction. Any other intermediates, including those from pBL do not interact with PBP. The reaction scheme and kinetics are determined. Interestingly, long-lived Complex-II dissociates upon excitation with blue light. These results demonstrate that Rc-PYP is a photochromic and new type of UV sensor, of which signaling process is similar to that of other light sensor proteins in the visible light region. The photochromic heterogeneous intermolecular interactions formed between PYP and PBP can be used as a novel and useful tool in optogenetics.

**Graphical Abstract:** 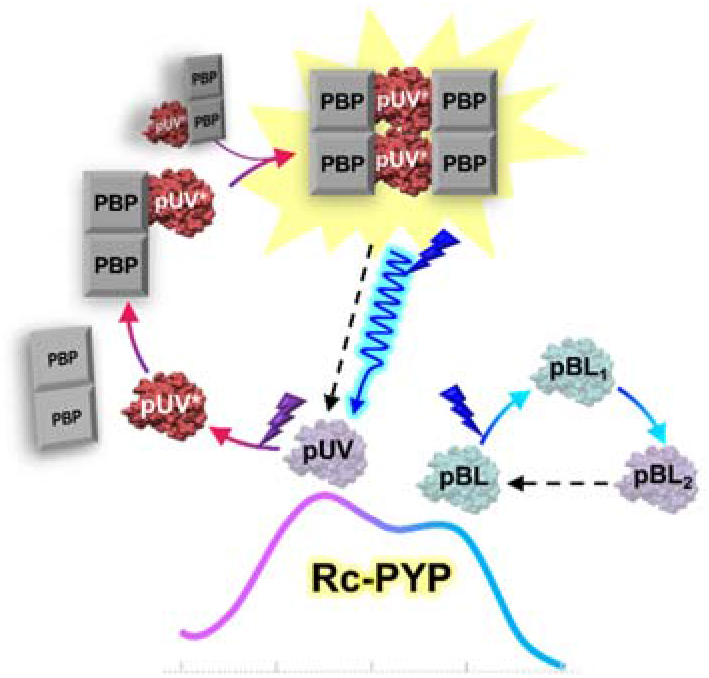

## Introduction

Photoactive yellow protein (PYP) is a light sensor protein that was initially found in the phototrophic bacterium *Halorhodospira halophila* (Hh-PYP) for a negative phototactic response.^1^ Since then, PYP has been used as a model protein to understand receptor activation in biological signal transduction processes. The reactions of Hh-PYP have been extensively studied using a variety of methods.^2–5^ The absorption spectrum of Hh-PYP in the near UV and visible wavelength region shows a maximum at 446 nm, and upon photoexcitation of this absorption band, the trans-cis isomerization of its chromophore, p-coumaric acid, occurs.^6^ After the isomerization, the red-shifted intermediate (pR), followed by the blue-shifted intermediate (pB) are generated.^7,8^ The pB species, which has a lifetime of ~1 s at physiological temperature, has been postulated to be the signaling state that transmits the light signal to downstream partner molecules.^3^ Hence, the biological function of Hh-PYP has been considered to be a blue light sensor.

Since PYP is one of the PAS domain proteins, that has various functions via intermolecular interactions,^9–11^ its downstream partner proteins have been of great interest. To date, downstream interaction sites have been predicted based on the sequence and three-dimensional structure compared with other PAS domain proteins.^12^ A biosensor based on diffusion detection has revealed that protein(s) from *Halorhodospira halophila* cell lysate interact with photoexcited Hh-PYP, indicating the presence of downstream molecules in the cell.^13^ However, even after a long time from the discovery of PYP, any downstream partner proteins have not been identified to date. In this study, we have investigated the intermolecular interaction dynamics of photoexcited PYP from *Rhodobacter capsulatus* (Rc-PYP) with a possible downstream partner protein, referred to as PYP-binding protein (PBP).

The photoreaction pathway for Rc-PYP has been previously reported.^14,15^ The absorption spectrum of Rc-PYP in the dark has two absorption maxima observed at 375 and 438 nm. These two bands represent two species, pUV in the ultraviolet (UV) light region and pBL in the blue light region.^15^ The different absorption wavelengths observed for pUV and pBL can be attributed to the different protonation states of the chromophore.^15^ Upon photoexcitation of pBL, isomerization of the chromophore takes place to form a cis-deprotonated species (pBL_1_) and subsequently, a protonated species (pBL_2_) is produced with a time constant of 82 μs.^15^ This pBL_2_ recovers back to the dark state with a lifetime of 1.2 ms.^15^ Upon photoexcitation of pUV, a cis-protonated intermediate species (pUV_1_) is formed, which undergoes structural changes including deprotonation with a time constant of 880 μs to form pUV_2_.^15^ A spectrally silent change in the conformation from this species with a time constant of 2.5 ms leads a long-lived form (pUV*).^15^ Thermal recovery to the dark state occurs with a lifetime of ~28 h.^15^ Comparing with the absorption spectrum obtained for Hh-PYP, we initially thought that one of intermediates of pBL could be a signaling state.

The PBP used in this study is a protein from a gene (RCAP_rcc01067) that is adjacent to PYP(RCAP_rcc01066) in the genomic sequence of *Rhodobacter capsulatus*.^16^ PBP is a 142-residue protein (SI-1). A secondary structural prediction of PBP using Jpred4 software predicts that PBP has α-helices at the N-terminus and β-sheets.^17^

We studied the interactions formed between Rc-PYP and PBP using size exclusion chromatography (SEC) and the kinetics by the transient absorption (TrA) and transient grating (TG) methods. This study clearly indicates that Rc-PYP is a unique photosensor protein, which is activated by UV light and negatively suppressed by blue light. Hence, Rc-PYP is a new type of UV sensor protein with photochromatic properties.

## Materials and methods

### Protein purification

Rc-PYP was produced and purified using a previously reported method.^15^ The DNA fragments encoding PBP from *R. capsulatus* were cloned into a pET-28a vector using a His-tag at the N-terminus and expressed in *E. coli* BL21 (DE3) cells. The *E. coli* strains were grown at 37 °C in LB medium supplemented with kanamycin (20 μg/mL) until the culture reached an OD_600_ of 0.6–0.8 and protein overexpression was induced upon the addition of isopropyl β-D-1-thiogalactopyranoside to a final concentration of 0.5 mM. The cells were incubated for 16–20 h at 25 °C. The harvested cells were suspended in 10 mM Tris-HCl, 100 mM NaCl, and 2 mM DTT at pH 8.0 and lysed upon ultrasonication. The protein was purified from the cell-free extract using an HisTrap HP column (GE Healthcare) and a linear imidazole gradient (30 to 500 mM) in 10 mM Tris-HCl, 100 mM NaCl, and 2 mM DTT at pH 8.0 to elute the protein from the HisTrap HP column. The His-tag was removed upon incubation of the target protein with thrombin (GE Healthcare) at room temperature for 16 h. For further purification, the samples were loaded onto an SEC column (Superdex 200 Increase 100/300 GL, GE Healthcare) using 10 mM Tris-HCl, 100 mM NaCl, and 2 mM DTT at pH 8.0. All measurements were performed in 10 mM Tris-HCl and 100 mM NaCl at pH 8.0, unless otherwise specified. DTT was removed immediately prior to each measurement. The concentration of PYP was calculated using the extinction coefficient at 375 nm (2.00 × 10^4^ M^−1^ cm^−1^), which was determined by the same method as previously reported.^18^ The concentration of PBP was determined using the extinction coefficient at 280 nm (1.55 × 10^4^ M^−1^ cm^−1^), which was calculated from the amino acid sequence by ProtParam.^19^

### Size exclusion chromatography (SEC)

SEC measurements were conducted using a Superdex 200 Increase 3.2/300 column (GE Healthcare) equilibrated with the Tris-HCl buffer at a flow rate of 0.05 mL min^−1^. The volume of the injected protein solution was 10 μL.

A Superdex 200 Increase 100/300 GL equilibrated with the Tris-HCl buffer was used to examine the composition of the Rc-PYP and PBP complex. The protein solution (100 μL) was injected and the flow rate was 0.4 min^−1^. The fraction around the target peak was collected 10 times and the solutions were concentrated to measure the absorption spectrum.

### Circular dichroism (CD) spectroscopy

The secondary structures of the Rc-PYP and PYP-PBP complexes were examined using CD spectroscopy (J-720WI, JASCO). A quartz cell with an optical path length of 0.2 cm was used. For measurement in the light state, UV light (360 ±5 nm) from an LED (NISSIN, UV3VP-8-365) passing through long- and short-wavelength pass filters was used. All measurements, including the CD and TG measurements were performed at 20 °C unless specified otherwise.

### TrA measurements

A laser pulse from a Nd:YAG laser (355 nm, SureliteII-10-T, Continuum) and a laser pulse from a Nd: YAG laser-pumped dye laser (480 nm, ND6000, Continuum) were used as the excitation light sources for our TrA measurements. Blue light at 450 nm from a diode laser (MicroLaser Systems) was used as the probe beam. The intensity of the probe beam was changed using a neutral-density filter. The probe beam was detected using a photomultiplier tube (R1477, Hamamatsu).

### TG measurements

The experimental set-up used for our TG experiments was similar to that previously reported.^20,21^ The excitation pulsed laser was the same as that used for our TrA measurements and a He-Ne laser (633 nm, JDS Uniphase) was used as the probe light source. The TG signal was detected using a photomultiplier tube. A quartz cell with an optical path length of 2 mm was used. The repetition rate of the pump pulse was 0.01 Hz. The sample solution was stirred after every excitation shot because the dark recovery of Rc-PYP was very slow. Typically, 30 signals were averaged using a digital oscilloscope (DSO9054H, Agilent). The grating wavenumber (*q*) was determined from the decay rate constant of the thermal grating signal obtained by a calorimetric reference sample.

### Analysis of TG signals

The principle of the TG method has been previously described.^22,23^ There are two dominant contributions to the TG signal under the present experimental conditions: the thermal grating due to the thermal energy from photoexcited molecules, and the species grating component due to the depletion of the reactant and the creation of intermediates and products. The TG signal intensity as a function of time (*I*_TG_(*t*)) is given by;

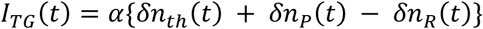

where α is a constant and δ*n*_th_(*t*)(<0) is the thermal grating component, which decays with a rate constant of *D*_th_*q*^2^ (*D*_th_: thermal diffusivity, *q*: grating wavenumber). The other terms, δ*n*_P_(*t*) and δ*n*_R_(*t*) are the species grating components; (δ*n*_spe_(*t*)) representing the changes in the refractive index due to the product and reactant, respectively. If the reaction completes faster than the observation time window and the molecular diffusion coefficient (*D*) is time-independent, the time profile of δ*n*_spe_(*t*) can be expressed as follows:

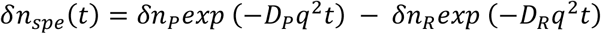

where *D*_R_ and *D*_P_ are the *D* values of the reactant and product, respectively. Furthermore, δ*n*_R_ (>0) and δ*n*_P_ (>0) are the initial changes in the refractive index due to the presence of the reactant and product, respectively. If *D* changes within the observation time window, the time profile should be analyzed considering this change.

## Results

### Detection of the PYP-PBP interaction using absorption spectroscopy

The PYP-PBP interaction was characterized by the absorption spectroscopy. Before showing the absorption spectrum of the Rc-PYP-PBP complex, the spectral changes associated with the Rc-PYP reaction without PBP are briefly described. As mentioned in the introduction, the absorption spectrum of Rc-PYP exhibits two peaks at 375 and 438 nm in the dark state. Upon photoexcitation by continuous wave (cw) UV light at 360 nm (UV(360)), the absorption band at 375 nm decreases and that at 438 nm increases (Fig. 1(a)). This spectral change was attributed to the formation of pUV* upon the photoexcitation of pUV. The lifetime of pUV* is very long (~28 h). Upon the photoexcitation of pBL by cw blue light at 480 nm (BL(480)), a similar change in the absorption spectrum was observed, but the change was very minor. This weak spectral change was attributed to the minor formation of pUV* in a very small yield.

**Fig. 1.**
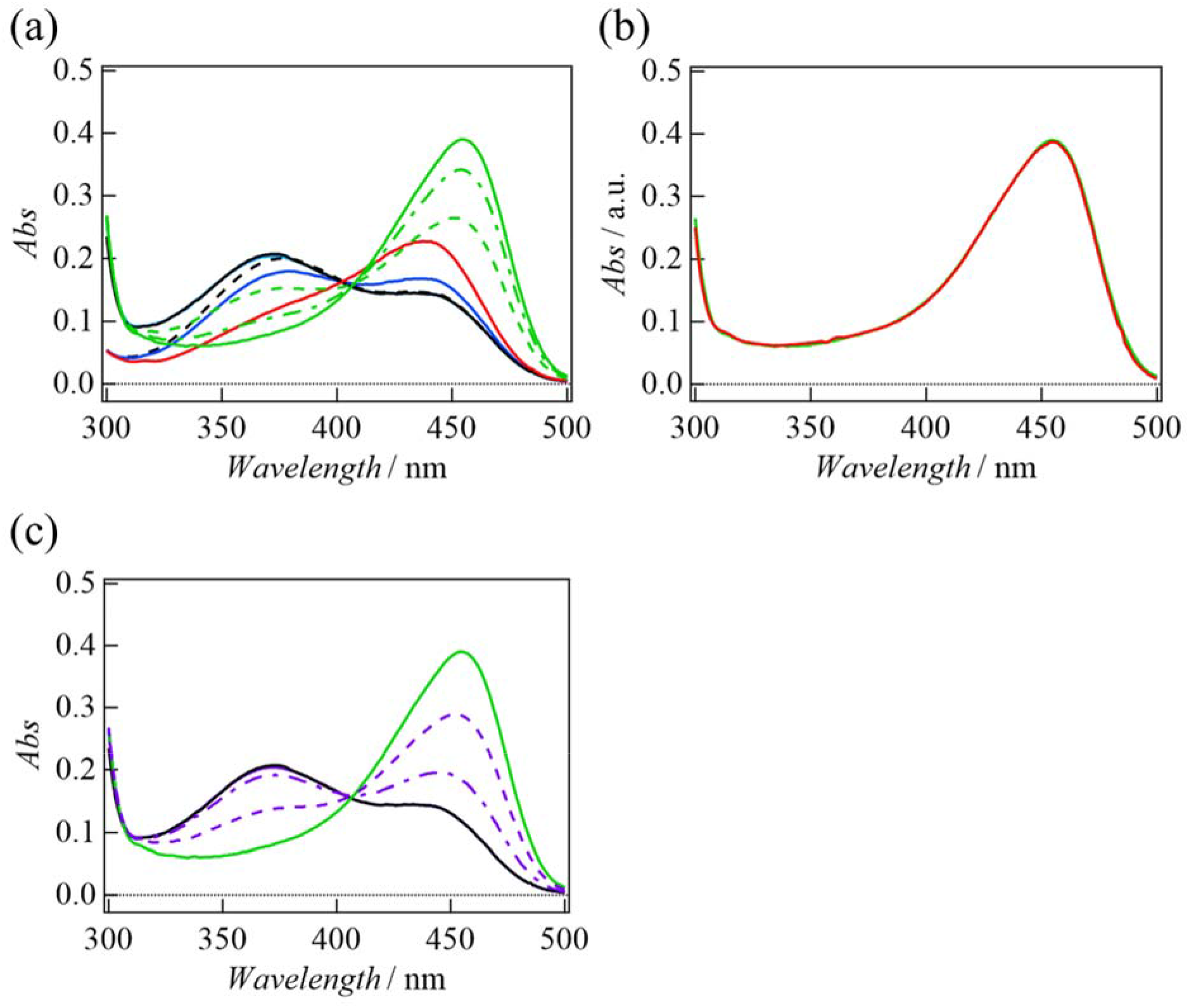
(a) Absorption spectra obtained for the PYP(10)-PBP(100) solution in the dark state (black solid line), after irradiation with 360 nm light for 1 s (green broken line) and 2 s (green dash-dotted line), and in the steady-state (green solid line). For comparison, the spectra obtained for the PYP solution (10 μM) in the dark state (black broken line), steady-state upon BL(480) irradiation with (blue solid line), and steady-state upon irradiation with UV(360) light (red solid line) are also shown. (b) The absorption spectra of the PYP(10)-PBP(100) solution irradiated with UV(360) light (red curve) and after mixing PBP to the pUV* solution (green curve) showing that both spectra are completely overlapped. (c) The absorption spectra obtained for the PYP(10)-PBP(100) solution in the dark state (black solid line) and in the steady state upon irradiation with UV(360) light (green solid line). The spectral changes upon BL(480) irradiation on the UV(360) pre-illuminated solution for 1 s (purple broken line) and 2 s (purple dash-dotted line), and in the steady state (purple solid line), which is completely overlapped by the black line, are also shown.

The absorption spectrum of a mixed solution at a Rc-PYP concentration ([Rc-PYP]) of 10 μM and PBP concentration ([PBP]) of 100 μM in the dark state is shown in Fig. 1(a). Hereafter, this solution prepared at [Rc-PYP] = 10 μM and [PBP] = 100 μM is referred as “PYP(10)-PBP(100) solution”. When compared with the absorption spectrum obtained for Rc-PYP without PBP, the absorption in a range of 320-380 nm is slightly enhanced. Since the absorption of PBP in this wavelength region is negligible, this change indicates that PYP somehow interacts with PBP in the dark state to influence the absorption spectrum. However, the interaction should be weak because this change is small. Indeed, the SEC measurements described below do not indicate a stable complex formed between Rc-PYP and PBP in the dark state. When the PYP(10)-PBP(100) solution was irradiated by BL(480) light, the absorption spectrum did not change. This behavior is different from that observed for Rc-PYP without PBP and explained in a later section. On the other hand, upon photoexcitation with UV(360) light, the absorption band at 375 nm gradually decreases and the absorption in the blue region (410–500 nm) significantly increases with an isosbestic point at 410 nm. In the steady state, the spectrum exhibits one peak at 455 nm. This spectrum is different from that obtained for Rc-PYP in the absence of PBP. Hence, this change is a clear indication of the interactions between PBP and excited Rc-PYP. The dark recovery of the spectrum is very slow and similar to that of Rc-PYP at least within 1740 min after the excitation (Fig. S2). A longer time trace of the recovery could not be measured due to aggregation. However, we speculate that the lifetime may be similar to that observed for pUV* (~28 h).

Previously, we have shown that two intermediates (pUV_1_ and pUV_2_) exist prior to forming the final stable pUV* product.^15^ To examine whether the PBP complex is formed with pUV* or one of the intermediates, we illuminated the Rc-PYP solution (10 μM) by UV(360) light to produce pUV* and then stopped the illumination to prepare a solution of pUV* without any intermediate species. PBP was added to the pUV* solution (final concentration of PBP: 100 μM) in the dark and the absorption spectrum was recorded (Fig. 1(b)). The absorption spectrum of the as-prepared solution was identical to that of the UV-irradiated PYP(10)-PBP(100) solution. Therefore, we conclude that PBP forms a complex with the stable pUV* product.

Interestingly, when the UV(360) pre-illuminated PYP(10)-PBP(100) solution is irradiated at 480 nm, the absorption spectrum gradually returns back to the spectrum obtained in the dark state (Fig. 1(c)). This observation suggests that the PYP-PBP complex is dissociated by the 480 nm light. Previously, we observed that the irradiation of pUV* with BL(480) results in the recovery of pUV.^15^ This blue light effect, recovery of pUV*, is consistent with the blue light-induced dissociation of the complex observed here.

### Oligomeric states of pUV*-PBP complex

The oligomeric states of Rc-PYP and PBP in the dark state were examined by the SEC method. The SEC profiles obtained for Rc-PYP at an injection concentration of 10 μM and PBP at 100 μM monitored at 280 nm exhibit a single peak at 13 and 36 kDa, respectively (Fig. 2(a)). Since the calculated mass of Rc-PYP is 14 kDa and that of PBP is 16 kDa, we conclude that Rc-PYP and PBP exist in their monomeric and dimeric forms, respectively.

**Fig. 2.**
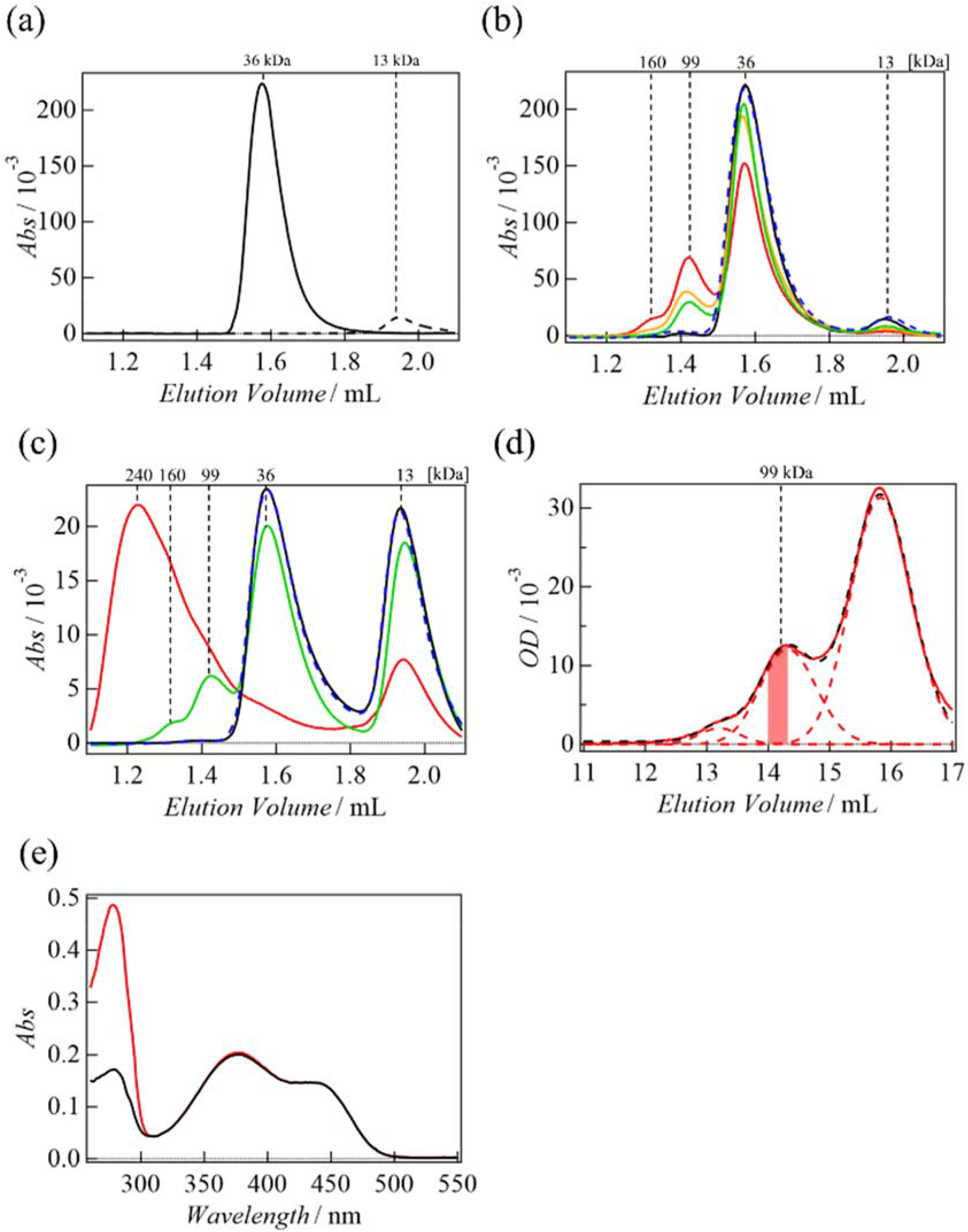
(a) SEC elution profile obtained for a solution of [PBP] = 100 μM (black solid line) and [PYP] = 10 μM (black broken line). (b) SEC profile obtained for the PYP(10)-PBP(100) solution in the dark state (black solid line), after UV(360) irradiation for 1s (green line) and 2 s (orange line), and in the steady state (red line). The SEC profile obtained upon BL(480) irradiation for the above steady state solution is also shown as a blue broken line, which is completely overlapped with the blue solid line. (c) The SEC profile obtained for a mixture of PBP at 10 μM with Rc-PYP at 10 μM in the dark state (black solid line), after UV(360) irradiation for 0.5 s (green solid line), and the steady state (red solid line). The SEC profile in the steady state of BL(480) illumination to the solution with pre-illuminated at 360 nm is also shown (blue broken line). (d) SEC profile obtained for the PYP(10)-PBP(100) solution monitored at 280 nm (red solid line) after UV(360) irradiation in the steady state. The best fitted curves of the profile by the lognormal functions are shown a black broken lines. The SEC profiles of the three components are separately shown by the red broken lines and the collected 99 kDa-fraction region is shown in red. (e) The absorption spectra of the 99 kDa component after the BL(480) irradiation (red curve) and the absorption spectrum of PYP in the dark state (black curve).

Next, the SEC profile of the PYP(10)-PBP(100) solution in the dark state was monitored at 280 nm. The profile is almost identical to the sum of those for the Rc-PYP and PBP solutions (Fig. 2(b)), although a very weak bump is observed at around 100 kDa. Hence, most of the Rc-PYP and PBP dimers exist separately in the dark state. Upon irradiation with UV(360) light for ~1 s, the intensities of the peaks observed at 13 and 36 kDa become weaker and a peak at 99 kDa appears. Upon increasing the irradiation period, the intensity of the peak at 99 kDa increases and a shoulder in higher molecular mass region appears. Hence, it is reasonable to propose that the photoexcited Rc-PYP (pUV*) initially creates a complex having a molecular mass of 99 kDa with PBP and larger complexes are formed upon the prolonged irradiation. When the UV-irradiated solution is illuminated at 480 nm, the profile is completely recovered to that observed in the dark state. This result supports the blue light dissociation pathway of this complex.

The SEC profile monitored at 280 nm for a mixed solution of [PYP] = 10 μM and [PBP] = 10 μM is shown in Fig. 2(c). Upon irradiation with UV light, the complex observed at 99 kDa is initially formed and subsequently, larger complexes of ~240 kDa are created. Since these larger complexes are not observed for the PYP(10)-PBP(100) solution even after prolonged irradiation, we propose that an excess amount of pUV* is required to produce the larger complexes. Upon BL(480) irradiation to this solution, the SEC profile of the solution recovers to that observed in the dark state. Hence, even the larger complexes dissociate back into Rc-PYP and the PBP dimer upon the blue light illumination.

The composition of the pUV*-PBP complex observed at 99 kDa was examined by isolating the SEC fraction as follows: The SEC profile of the UV(360)-irradiated PYP(10)-PBP(100) solution is decomposed into three components by fitting with three lognormal functions (Fig. 2(d)). The main band fraction of the 99 kDa component without contamination by the adjacent 160 and 36 kDa components (the colored region in Fig.2(d)) is collected. This solution is irradiated at 480 nm to recover the dark state and the absorption spectrum is measured (Fig. 2(e)). This spectrum should be the sum of the Rc-PYP and PBP spectra. The spectrum is fitted by the spectrum obtained for Rc-PYP (Fig. 1) in the visible wavelength region (350–500 nm), as shown in Fig. 2(e). The ratio of the absorbance of PBP (A_PBP_) to PYP (A_PYP_) at 280 nm is determined from these spectra to be A_PBP_/A_PYP_ = 1.79. Using the absorption coefficients of PBP and PYP at 280 nm (1.55 × 10^4^ M^−1^ cm^−1^ and 1.71 × 10^4^ M^−1^ cm^−1^), the molar ratio of PBP to PYP is calculated to be 1.97. Since the molecular mass of this complex is 99 kDa, we identify that this complex consists of four PBP (two dimers) and two pUV* molecules, i.e. pUV*_2_-PBP_4_ (calculated molecular mass: 92 kDa).

In the following sections, the dynamics of light-dependent complex formation for the PYP(10)-PBP(100) solution with a weak pulsed laser light, under which conditions pUV*_2_-PBP_4_ is formed, are studied by the TrA and TG measurements.

### Dynamics of complex formation monitored by TrA

The reaction kinetics upon pulsed UV light irradiation at 355 nm were studied using the TrA method. Fig. 3(a) shows the TrA signal after excitation at 355 nm and probed at 450 nm for the Rc-PYP and PYP(10)-PBP(100) solutions. The TrA signal corresponding to Rc-PYP without PBP has been previously reported. According to this result, the pUV_1_ intermediate is formed after the UV excitation and the TrA signal changes single-exponentially with a time constant of 880 μs, indicating the formation of the pUV_2_ intermediate. After this process, the final pUV* product is formed via a spectrally silent process with a time constant of 2.5 ms. Upon adding PBP to this solution, a slower component appears in addition to the signal corresponding to Rc-PYP. Since PBP interacts with the stable pUV* product, as demonstrated in the above section, this slower component should represent the time course for the formation of the PBP-pUV* complex. This time constant does not depend on the excitation light intensity. If this component represents the formation of pUV*_2_-PBP_4_, the time profile should exhibit the second-order kinetics with respect to [pUV*] and should depend on the excitation light intensity. The light intensity independence suggests that this component is associated with PBP2 with one pUV* (Complex-I). We consider that pUV*_2_-PBP_4_ (Complex-II) is formed via the dimerization of Complex-I. This suggestion is supported by the PBP concentration dependence and excitation light intensity dependence of the TG signals, as described below.

**Fig. 3.**
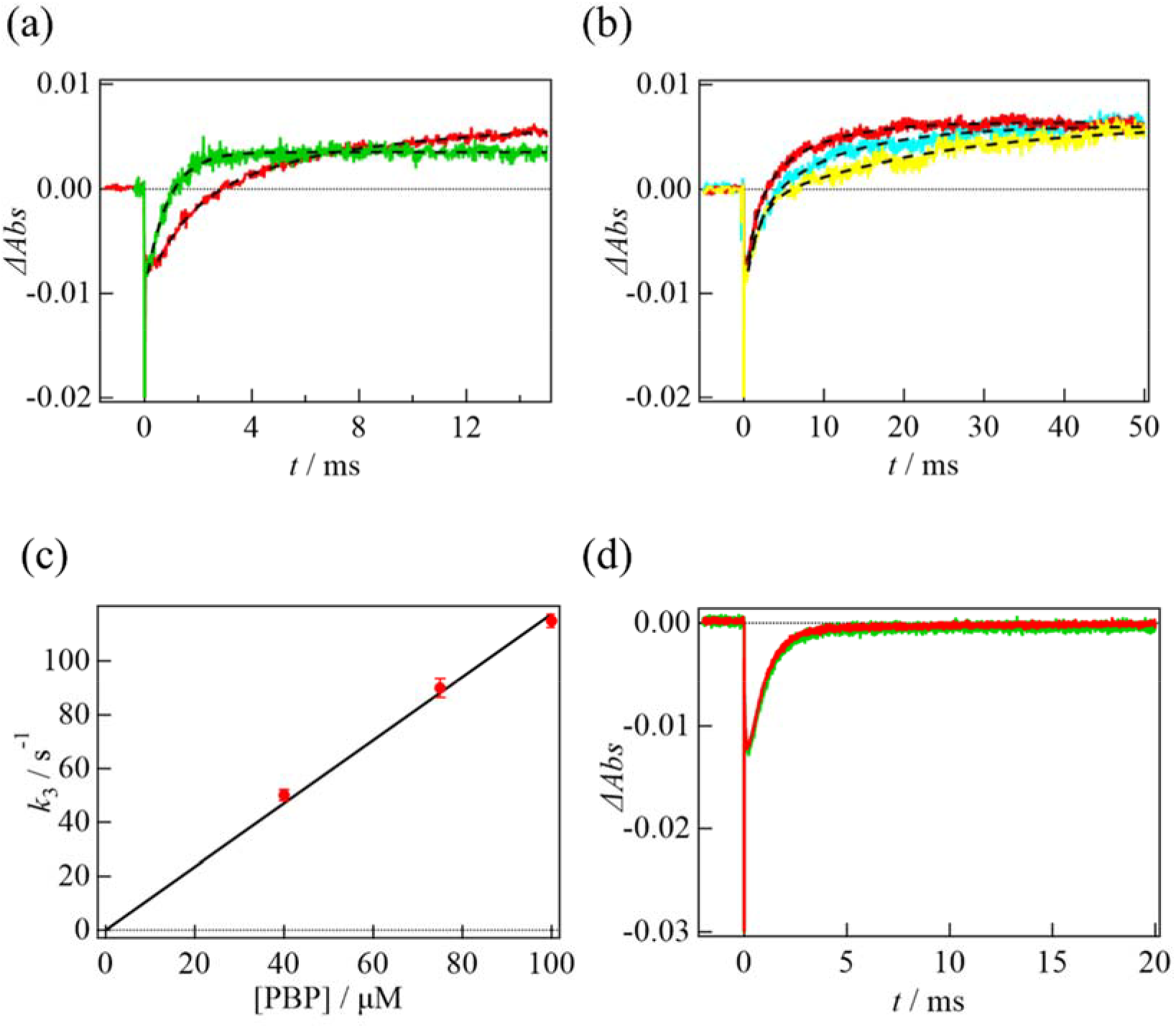
(a) The TrA signal obtained for the PYP(10)-PBP(100) solution (red curve) and the solution at [Rc-PYP] = 10 μM in the absence of PBP (green curve) upon excitation at 355 nm and probed at 450 nm. The best fitted curves by a single-exponential function for the Rc-PYP solution and by eq.(1) for the PYP(10)-PBP(100) solution are shown by the black broken lines. (b) The PBP concentration dependence of the TrA signal at [PYP] = 10 μM upon excitation at 355 nm and probed at 450 nm. The PBP concentrations are 100 μM (red), 75 μM (blue), and 40 μM (yellow). The best fitted curves by eq.(1) are shown by the black broken lines. (c) The plot of *k*_3_ determined from the TrA signals vs [PBP]. The black line is the best fitted line with a linear function. (d) A comparison of the TrA signal obtained for the PYP(10)-PBP(100) solution (red curve) and Rc-PYP without PBP (green curve) upon excitation at 480 nm and probed at 450 nm.

Based on the above consideration, we analyzed the TrA time profile based on Scheme 1:

**(Scheme 1).**
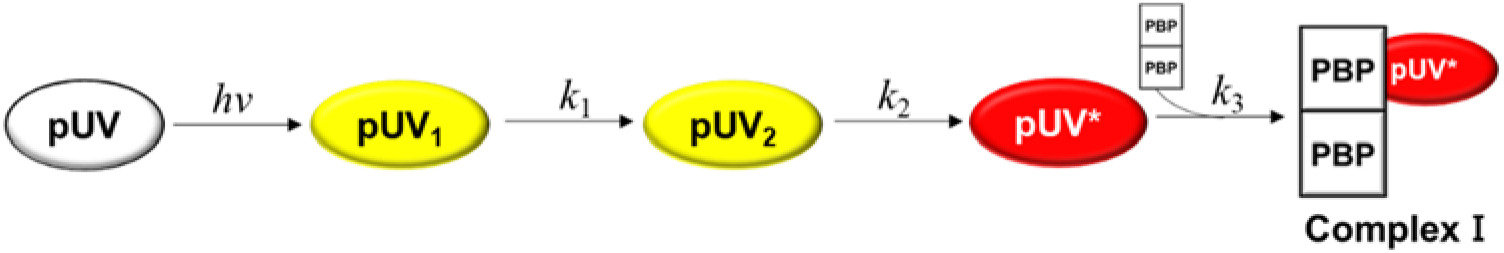

where *k*_i_ (i = 1–3) is the rate constant. We assume that the rate of the last step is the pseudo-first-order reaction due to the sufficiently high concentration of PBP. The time profile of the TrA signal (I_TrA_(t)) based on Scheme 1 should be expressed by a sum of three exponential functions as follows:

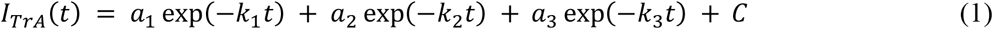

where a_i_ (i = 1–3) are pre-exponential factors and C is a constant. To fit the signal by this equation, we fixed the rate constants at *k*_1_ = (880 μs)^−1^ and *k*_2_ = (2.5 ms)^−1^, which were determined previously, to reduce the ambiguities of the fitting. The signal is well reproduced by this equation and *k*_3_ is determined to be (8.6 ±0.5 ms)^−1^ at [PBP] = 100 μM.

We found that the rate constant *k*_3_ depends on [PBP] (Fig. 3(b)). The *k*_3_ values determined at several PBP concentrations are plotted against [PBP] in Fig. 3(c). From the slope of the fitting, the second-order rate constant for the Complex-I formation is determined to be 1.2 (±0.03) × 10^6^ M^−1^s^−1^.

In the above section, we have shown that PBP does not form a complex upon the pBL excitation. However, it is possible that PBP forms a complex with one of the intermediates during the pBL reaction (pBL_1_ and pBL_2_) and returns to the dark state via a fast recovery process. To examine this possibility, we measured the TrA signal of Rc-PYP in the presence and absence of PBP upon pulsed-BL(480) excitation and probed at 450 nm (Fig. 3(d)). The profile of the signal does not depend on the presence of PBP at all suggesting that the intermediates produced by photoexcitation of pBL do not interact with PBP.

### Association reaction detected by TG method

The TG signals of Rc-PYP at 200 μM and the PYP(10)-PBP(100) solution upon pulsed-UV excitation at 355 nm and *q*^2^ = 3.8 × 10^10^ m^−2^ are shown in Fig. 4(a). We previously reported the TG signals of Rc-PYP and their analysis.^15^ The thermal grating signal rises within 20 ns, decays to the baseline, and shows a very weak molecular diffusion signal for the Rc-PYP solution in the absence of PBP. Upon adding PBP to the solution, the thermal grating signal does not change, but a very strong rise-decay signal appears in the 10–1000 ms time range at this *q*^2^. Since the time scale of this component depends on *q*^2^ (Fig.4(b)), this component is assigned to a protein diffusion signal. Considering that the change in the refractive index of the thermal diffusion signal is negative, the signs of the change in the refractive index for the rise and decay components of the diffusion signal are determined to be the negative and positive, respectively. Therefore, the rates of the rise and decay components represent the diffusion of the reactant and product, respectively, i.e. *D* decreases upon photoexcitation. The decrease in *D* is reasonable for the association reaction between pUV* and PBP.

**Fig. 4.**
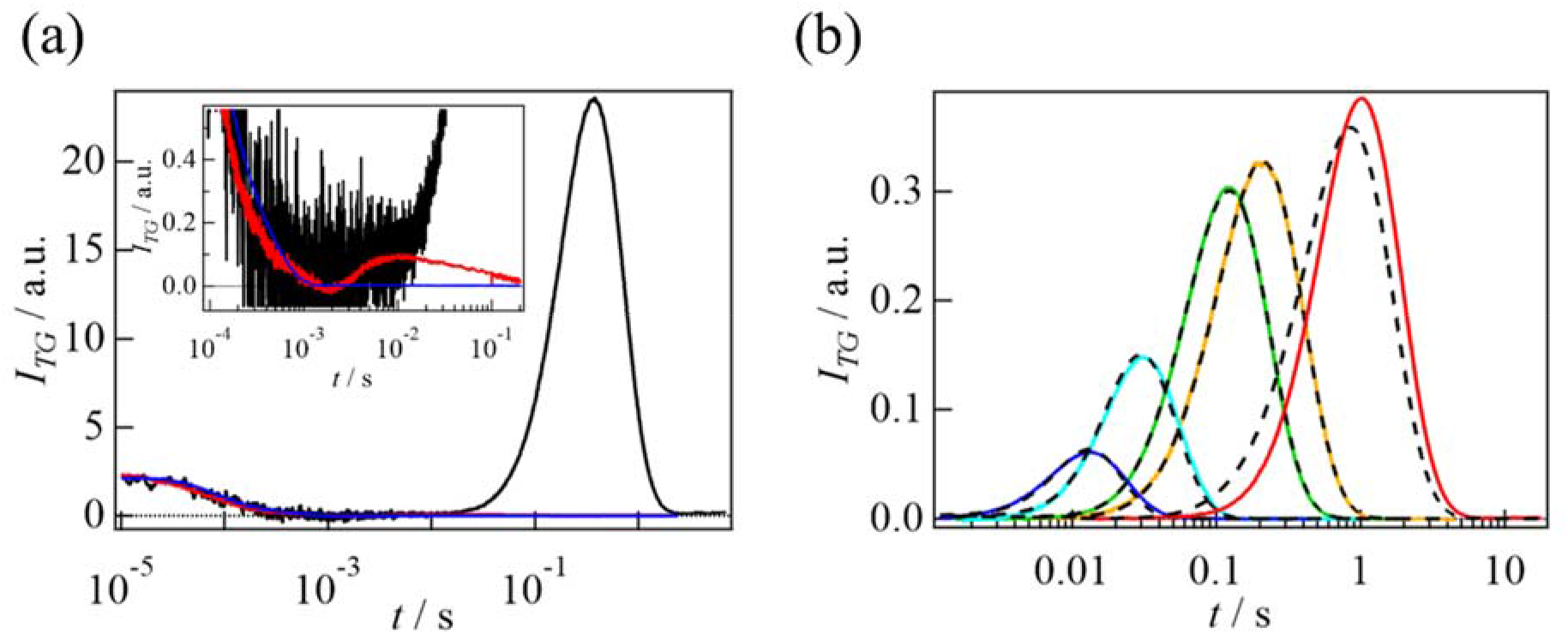
(a) The TG signals of Rc-PYP (red), the PYP(10)-PBP(100) solution upon excitation at 355 nm (black), and the PYP(10)-PBP(100) solution upon excitation (blue) at 480 nm measured at *q*^2^ = 3.8 × 10^10^ m^−2^. The magnified signals in a fast time region are shown in the inset. (b) The *q*^2^ dependence of the TG signal of the PYP(10)-PBP(100) solution. The *q*^2^ values are 1.3 × 10^12^, 5.4 × 10^11^, 1.1 × 10^11^, 6.3 × 10^10^, and 1.0 × 10^10^ m^−2^ from left to right. The best-fitted curves by eq.(S1) in the range of *q*^2^<6.3 × 10^10^ m^−2^ are shown as black broken lines.

The kinetics of the *D*-change is investigated by the TG method at various *q*^2^ values (Fig. 4(b)). The signal intensities are normalized by the intensity of the thermal grating signal measured under the same experimental conditions, which is proportional to the square of the number of the photoexcited molecules. The diffusion signal intensity gradually increases with a decrease in *q*^2^. This behavior indicates that *D* gradually decreases in this time range.

We also examined the excitation light intensity dependence at various *q*^2^. After the normalization by the thermal grating signal from the calorimetric reference sample, the signal does not depend on the light intensity in a range of q^2^>3.1×10^10^ m^−2^ (Fig.5(a)). However, at a small *q*^2^, such as *q*^2^ = 1.0 × 10^10^ m^−2^, the intensity of the diffusion signal increases with increasing the light intensity (Fig.5(b)). Moreover, the peak of the signal becomes faster with increasing the light intensity (Fig.5(c)). These observations suggest that the diffusion signal at this small *q*^2^ comes from the bimolecular reaction of two photoexcited species. Hence, the association reaction between pUV* and PBP_2_ occurs first to produce Complex-I in the time range of *t* <1 s at this light intensity and subsequently, two Complex-I molecules associate to produce pUV*_2_-PBP_4_ (Complex-II).

**Fig. 5.**
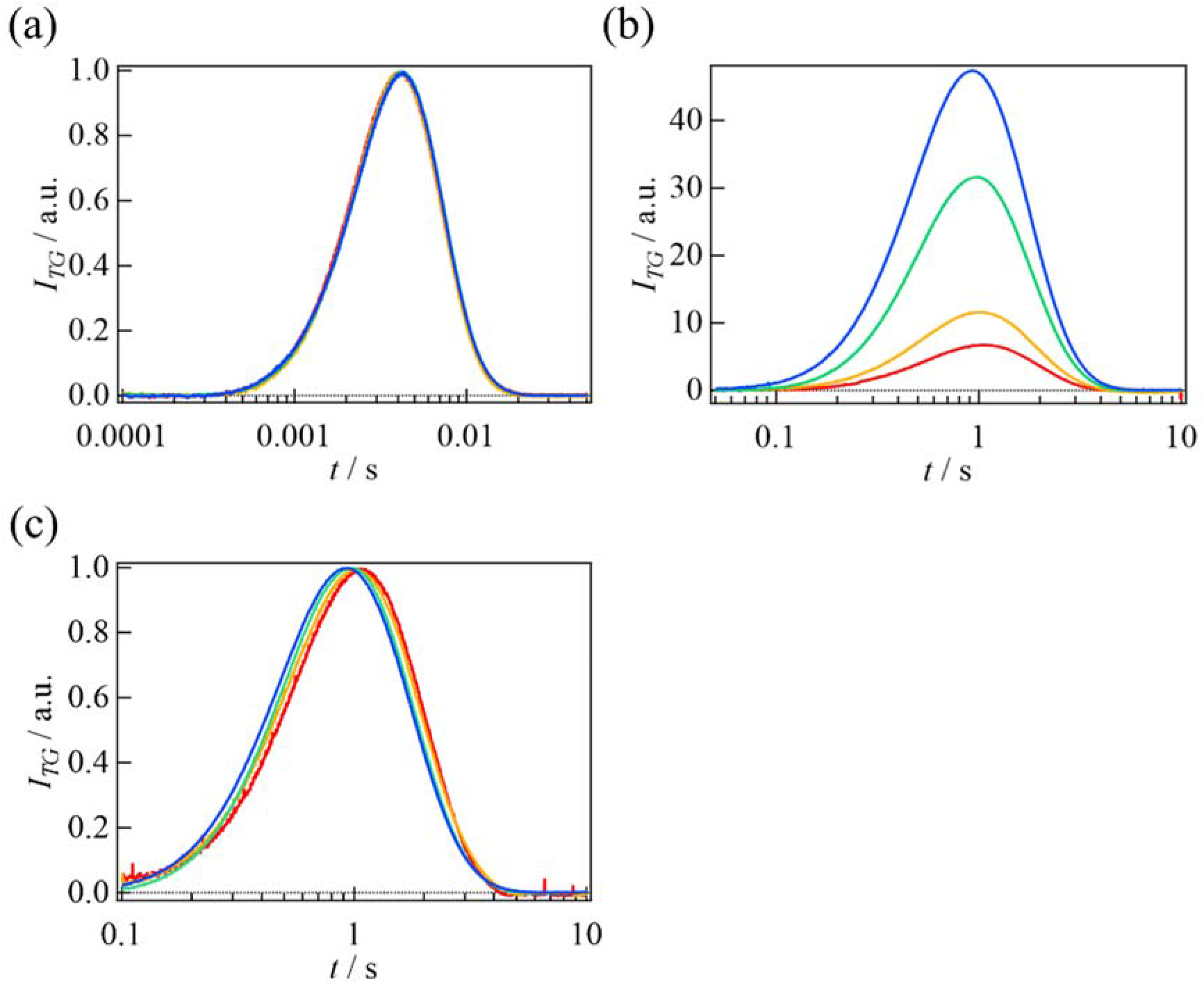
Laser power dependence of the TG signal of the PYP(10)-PBP(100) solution measured at *q*^2^ = (a) 5.0 × 10^12^ and (b) 1.0 × 10^10^ m^−2^. The laser powers are 6.7 (red), 11 (orange), 29 (green), and 55 (blue) J/m^2^. (c) The diffusion signals of (b) are normalized at the peak intensity.

The *q*^2^-dependent signals that do not depend on the excitation light intensity are globally analyzed based on Scheme 1. Here, since the rate of the pUV_2_ creation (*k*_1_-phase) is faster than the observation time window of the diffusion signal, we assume that pUV_2_ is created sufficiently fast upon the pulse excitation. Using this model, an analytical equation describing the diffusion signal is derived and presented in eq.(S1). To fit the signals using eq.(S1), *k*_2_ and *k*_3_ are fixed with the values obtained from our TrA measurements. Furthermore, the diffusion coefficients for pUV (*D*_UV_), pUV_2_ (*D*_UV2_), and pUV* (*D*_UV*_) are fixed to those obtained in the previous study.^15^ Hence, the adjustable parameters for the fitting are the diffusion coefficients of Complex-I (*D*_C1_), PBP (*D*_PBP_), and the change in the refractive index. This calculated signal agrees well with the observed signals and the determined values are listed in Table 1.

**Table 1.** Reaction properties upon excitation at 355 nm determined by TG signals.

| Diffusion coefficient ($10^{-11}$ m<sup>2</sup>/s) | | | | | Rate constant | |
| --- | --- | --- | --- | --- | --- | --- |
| $D_{UV}$ | $D_{UV2}$ | $D_{UV*}$ | $D_{C1}$ | $D_{PBP}$ | $k_2$ | $k_3$ |
| 12 | 8.5 | 12 | $6.1 \pm 0.3$ | $9.8 \pm 0.4$ | $(2.5 \text{ ms})^{-1}$ | $(8.6 \text{ ms})^{-1}$ |

The dependence of the TG signal on the concentration of PBP is shown in Fig. 6(a). After normalization by the thermal grating signal intensity, the diffusion signal increases with increasing [PBP], and shifts faster (Fig. 6(b)), indicating that the rate of Complex-I formation increases. The observed signals are analyzed by eq.(S1) with a restriction that only *k*_3_ is the adjustable parameter. The determined *k*_3_ value is proportional to [PBP] (Fig. 6(c)) and the second order rate constant is determined to be 1.1 (±0.04) × 10^6^ M^−1^s^−1^. This value is in good agreement with that determined using the TrA signal. This indicates that the slow rise observed in the TrA signal represents the formation of Complex-I.

**Fig. 6.**
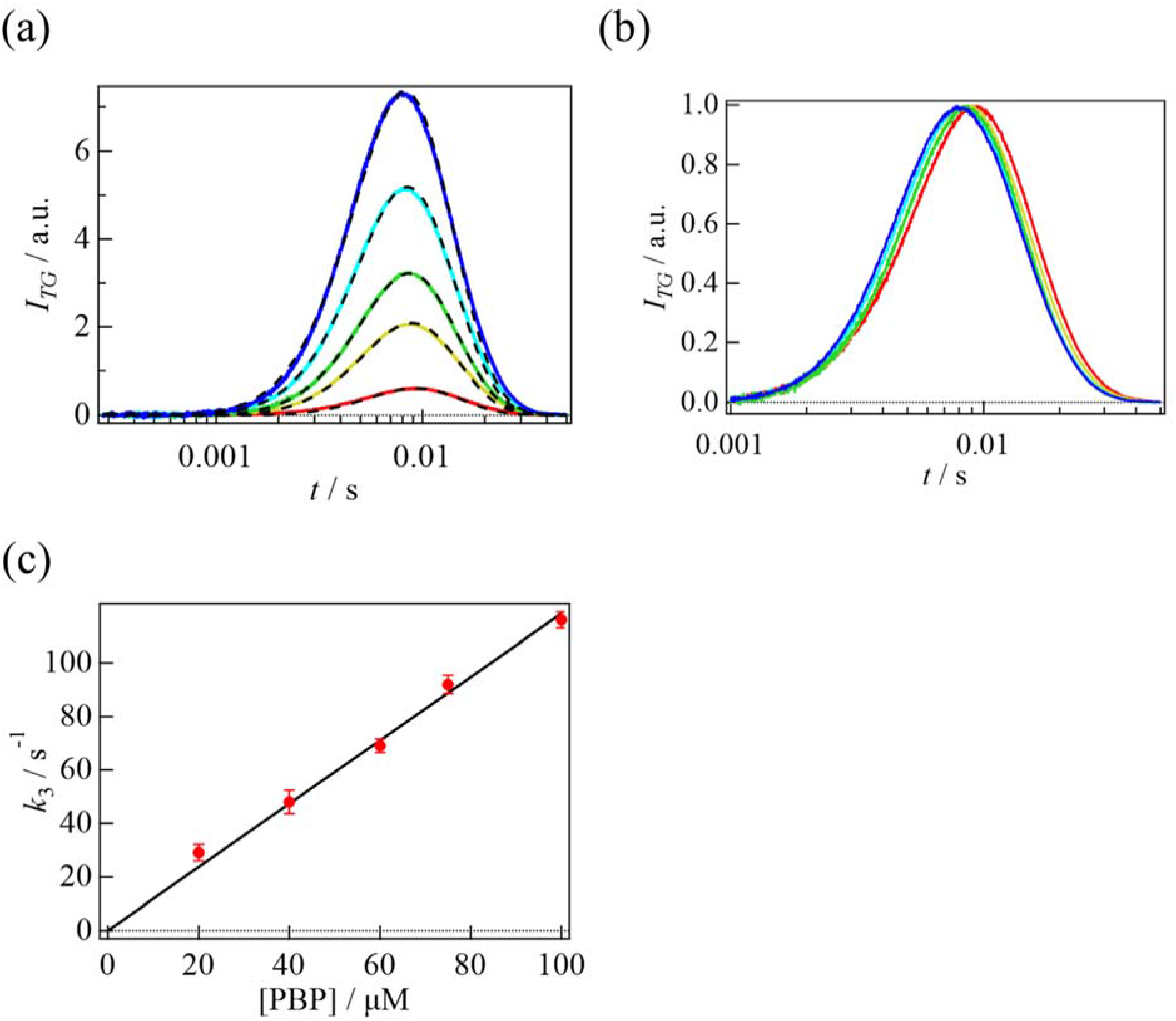
(a) The PBP concentration dependence of the TG signal measured at *q*^2^ = 2.2 × 10^12^ m^−2^. The concentration of PYP is 10 μM and the concentration of PBP is 100 μM (blue), 75 μM (pale blue), 60 μM (green), 40 μM (yellow), and 20 μM (red). The best-fitted curves based on eq.(S1) are shown using black broken lines. (b) The diffusion signals of (a) are normalized at the peak intensity. (c) Plot of *k*_3_ determined from the TG signal vs [PBP]. The black line is the best fitted line using a linear function.

According to the Stokes–Einstein relationship, *D* is inversely proportional to the radius of the protein. Assuming that all species are spherical in shape, the radius of Complex-I is larger than that of Rc-PYP by a factor of (46 kDa/14 kDa)^1/3^ ~1.49. Hence, the *D* of Complex-I is calculated to be 8.0 × 10^−11^ m^2^/s, which is much larger than that observed (6.1 × 10^−11^ m^2^/s). If the shape of the molecule deviates significantly from a sphere, the *D* value may be different from that calculated above. However, the deviation from the sphere may not be large, because the molecular mass determined using SEC is in good agreement with the calculated mass. Instead, the origin of the discrepancy between the predicted and observed *D* values may be due to conformation changes that cause an increase in the friction for translational diffusion. Indeed, previous studies on many photosensor proteins have indicated that the unfolding of the secondary structure causes a decrease in *D*.^24–26^ We measured the CD spectra of Rc-PYP in the presence and absence of PBP in the dark state and upon irradiation at 360 nm (Fig. 7(a)). In the case of Rc-PYP in the absence of PBP, the change in the CD spectrum is very small, but in the presence of PBP, a slight change in the CD spectrum is observed at ~208 nm (Fig. 7(a)). Based on this result, we suggest that the significant decrease in *D* due to the formation of Complex-I is due to the secondary structure change caused by the association between pUV* and PBP.

**Fig. 7.**
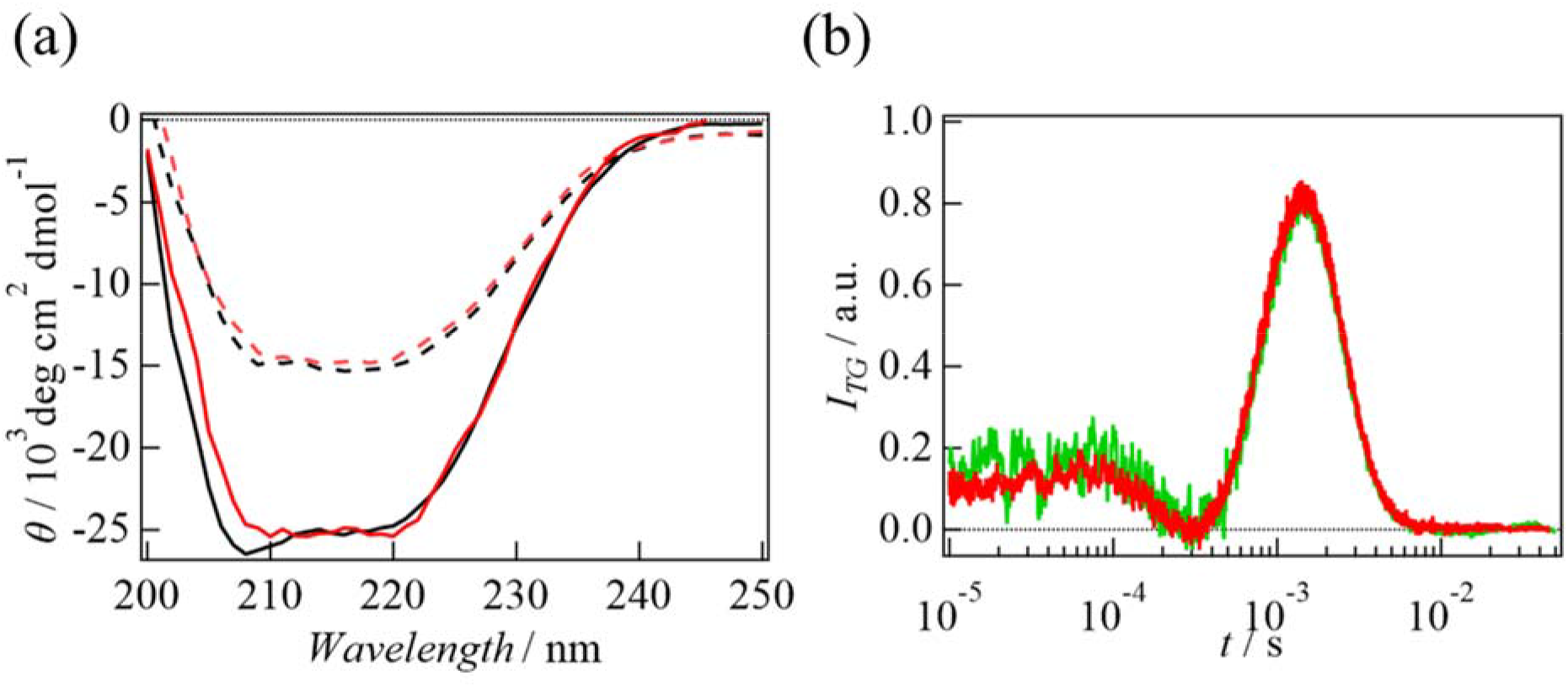
(a) CD spectra of a solution of [Rc-PYP] = 1 μM and [PBP] = 10 μM (black solid line), and after irradiation at 360 nm (red solid line). For comparison, the CD spectra of Rc-PYP at 10 μM in the dark state (black dotted line) and after irradiation at 360 nm (red dotted line) are also shown. (b) The TG signals of Rc-PYP at 50 μM in the absence (red) and presence of PBP (500 μM) (green) upon excitation at 480 nm.

In the above section, we have suggested that PBP does not interact with any intermediates during the pBL reaction based on our TrA measurements. When we compare the TG signal of PYP(10)-PBP(100) excited at 480 nm with that at 355 nm under the same conditions, the intensity is negligibly weak (Fig. 4(a)). To further confirm this, the interaction between PBP and the intermediates in the pBL reaction is examined by the TG method in a shorter time range. If the interaction between PBP and the intermediates of the pBL reaction changes, the change should manifest as a diffusion change. The TG signals of PYP at 50 μM and the PYP(50)-PBP(500) solution upon excitation at 480 nm under the same conditions are shown in Fig.7(b). No change in the TG profile in the time range of 0.01-10 ms is observed, indicating that the intermediates produced by photoexcitation of pBL do not interact with PBP.

It is clear that the TG signal of the smallest *q*^2^ value cannot be analyzed by eq.(S1) (Fig. 4(b)) and the excitation light intensity dependence is observed only at this smallest *q*^2^ value (Fig. 5(b)). This light intensity dependence can be explained by the second-order reaction rate of the Complex-II formation on the excited molecules (Complex-I). Because the time range for Complex-II formation is too slow to be analyzed, it is not possible to determine the diffusion coefficient and formation rate of Complex-II. However, from the time scale of the light intensity-dependent molecular diffusion signal, we consider that the formation of Complex-II occurs within a few seconds. The TrA signal in the presence of PBP does not show any component slower than the absorption change for Complex-I formation, suggesting that the reaction of Complex-I to form Complex-II is not accompanied by a change in absorption.

### Dissociation process induced by blue light

We show that Complex-II dissociates upon BL(480) irradiation in the previous sections. This dissociation reaction process is studied by the TrA and TG methods. The Complex-II solution is prepared by the UV(360) irradiation to the PYP(10)-PBP(100) solution. Fig. 8(a) shows the TrA signal of this UV-pre-illuminated solution upon pulse-excitation at 480 nm and probed at 450 nm. The profile consists of fast and slow decay phases. For comparison, the TrA signal detected by the same procedure in the absence of PBP is also shown in Fig. 8(a). This signal represents the reverse reaction from pUV*, which can be fitted well by a single exponential function with a time constant of 810 ±50 μs in this time range and sensitivity. The time profile of the TrA signal of the Complex-II solution is well reproduced by a biexponential function with time constants of 820 ±15 μs and 10 ±2 ms. The faster time constant of 820 μs is almost identical to that observed in the Rc-PYP solution. Therefore, we attribute this 820 μs-component observed in the PYP(10)-PBP(100) solution to the recovery kinetics from pUV* to pUV in Complex-II before dissociation. The almost identical values of the rate constants observed for pUV* and Complex-II indicate that the recovery reaction of pUV* is not influenced by the formation of the complex with PBP. Since the phase at 10 ms is not observed in the complex formation process and the rate does not depend on the concentration of PBP (Fig. 8(b)) or the excitation light intensity, this component is attributed to the dissociation process of the PYP-PBP complexes.

**Fig. 8.**
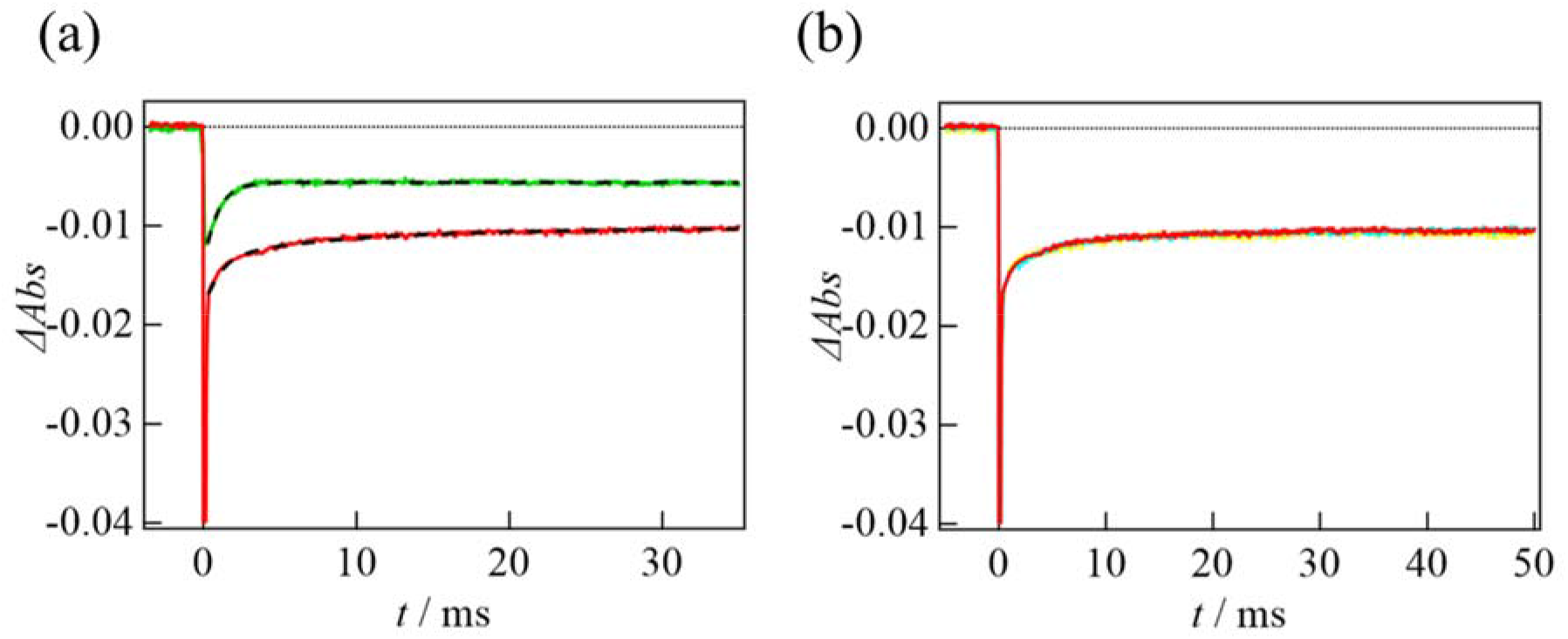
(a) TrA signal of the UV pre-illuminated Rc-PYP solution (10 μM) (green curve) and the PYP(10)-PBP(100) solution (red curve) upon excitation at 480 nm and probed at 450 nm. The best fitted curves by the single-exponential and bi-exponential functions for these signals are shown by the broken lines. (b) The PBP concentration dependence of the TrA signal for the UV pre-illuminated solution probed at 450 nm. The concentration of PYP is 10 μM and concentration of PBP are 100 (red), 75 (blue), 40 μM (yellow).

Fig. 9(a) shows the TG signal excited at 480 nm for the UV(360) pre-illuminated PYP(10)-PBP(100) solution at *q*^2^ =3.9 × 10^10^ m^−2^. The thermal grating signal decays via a thermal diffusion process and a rise-decay profile is observed. Because the rise-decay signal depends on *q*^2^ (Fig. 9(b)), this component is the diffusion signal. It should be noted that the thermal grating signal does not decay to the baseline, indicating that the signs of *δn* for the rise and decay components are positive and negative, respectively. Hence, the rise and decay rates are attributed to the diffusion of the product and reactant, respectively, i.e. the product diffuses faster than the reactant, which indicates the dissociation reaction.

**Fig. 9.**
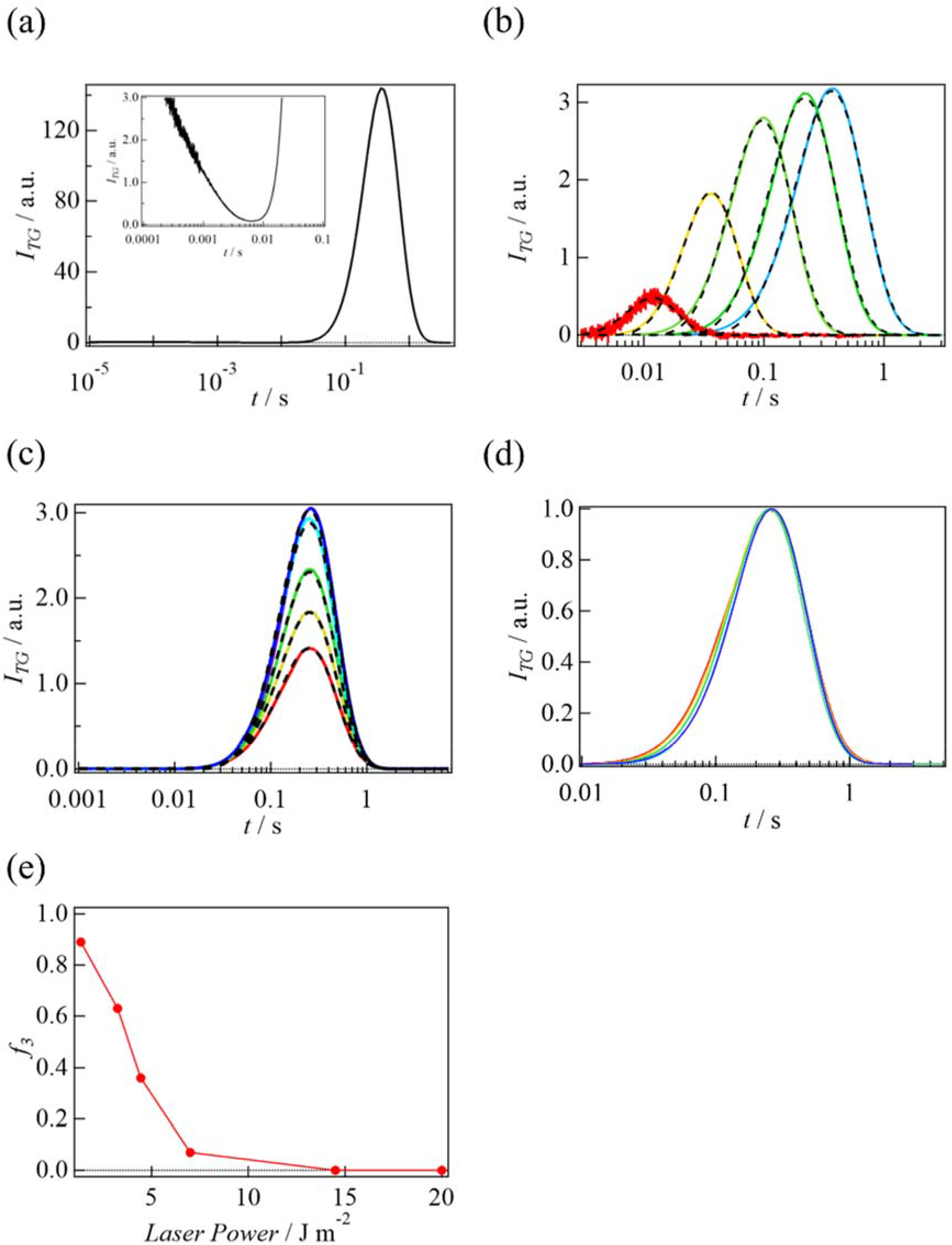
(a) The TG signal of the UV light pre-illuminated PYP(10)-PBP(100) solution upon excitation at 480 nm measured at *q*^2^ = 3.9 × 10^10^ m^−2^. The magnified signal in a fast time region is shown in the inset. (b) The *q*^2^ dependence of the TG signal of the UV pre-illuminated PYP(10)-PBP(100) solution upon excitation at 480 nm. The *q*^2^ values are 2.3 × 10^12^, 6.1 × 10^11^, 1.7 × 10^11^, 7.2 × 10^10^, and 3.9 × 10^10^ m^−2^ from left to right. The best-fitted curves based on Scheme 2 (eq.(S2)) are shown using black broken lines. (c) The laser power dependence of the diffusion signal of the UV(360) pre-illuminated PYP(10)-PBP(100) solution measured at *q*^2^ = 5.7 × 10^10^ m^−2^, normalized by the species grating signal intensity observed before the diffusion signal. The laser powers are 1.4 (red), 3.3 (yellow), 4.5 (green), 7.0 (cyan), 15 (blue), and 20 (purple) J/m^2^. The best-fitted curves based on eq.(S2) are shown using black broken lines. (d) The laser power dependence of the diffusion signal normalized at peak intensity observed under the same conditions as (c). (e) Plot of the relative contribution of Scheme 3 (f_3_) vs the laser power.

The dissociation reaction kinetics is studied by the *q*^2^-dependence. The diffusion signals excited at 480 nm for the UV(360) pre-illuminated PYP(10)-PBP(100) solution at various *q*^2^ values are shown in Fig. 9(b). The signals are normalized by the species grating signal intensity observed before the diffusion signal, which represents the number of photo-reacted species. These signals are measured at a sufficiently strong excitation light intensity, at which the profile of the signal does not depend on the light intensity. At this excitation light intensity, we consider that two pUV* molecules are excited in Complex-II, which dissociates into two pUV molecules and two PBP dimers (Scheme 2), as confirmed later. The analytical equation for the time dependence of the species grating based on this pathway (δn_S2_) is given by eq.(S2). For the global fitting of the signals, *D*_UV_ and *D*_PBP_ are fixed at the values obtained from the measurement of the association reaction dynamics described in the above section. Here, since the rate constants of the back reaction of pUV* (*k*_UV*_) is faster than the observation time window for the diffusion signal, the rate of this process is neglected for the analysis. Moreover, the rate constant of the dissociation (*k*_dis_) is fixed at the values obtained from the TrA analysis and we assumed that the diffusion coefficients of Complex-II (*D*_C2_) are the same as those of the intermediate (pUV_2_-PBP_4_). Hence, the adjustable parameters are the refractive index changes, and *D*_C2_. Even under these restrictions, the observed signals are reproduced well by this equation and the determined parameters are listed in Table 2.

**Scheme 2.**
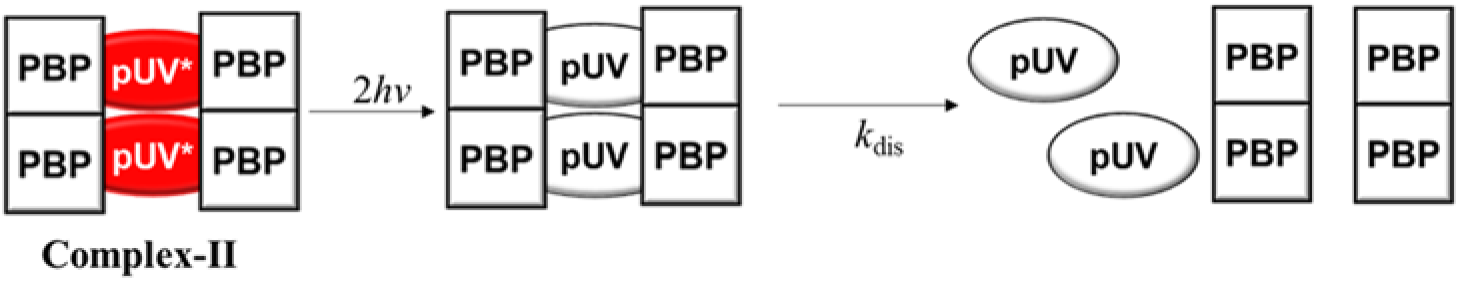

**Table 2.**
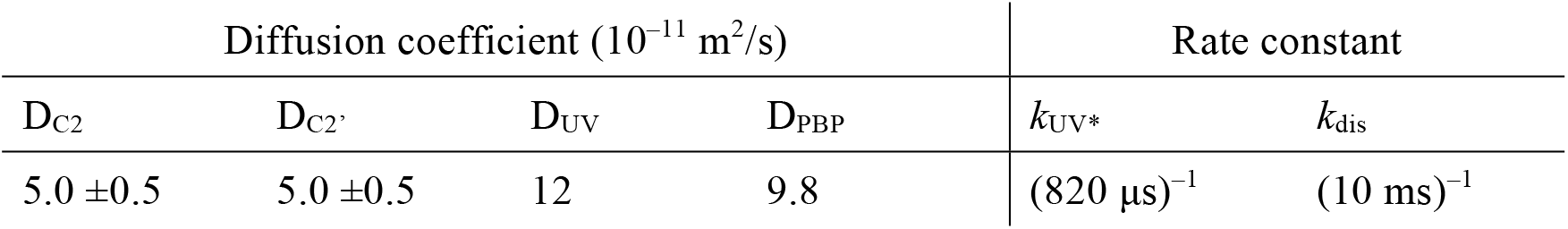
Reaction properties of the TG signal upon excitation at 480 nm after UV pre-illumination.

The diffusion signals measured at various excitation light intensities are shown in Fig. 9(c). The signal intensity increases as the excitation light intensity increases and becomes saturated above 15 J/m^2^. It is interesting to note that the profile slightly depends on the light intensity in the weak light region (Fig. 9(d)). Although the difference is small, this behavior is reproducible. This light intensity dependence indicates that the reaction scheme upon the excitation of one pUV* molecule in Complex-II is different from that upon the excitation of two pUV*. It may be reasonable to consider that when one pUV* molecule is excited, Complex-II dissociates into pUV, PBP dimer, and Complex-I (Scheme 3). The analytical equation based on this pathway (δn_S3_) is given by eq.(S3). It is apparent that the calculated signals based on Scheme 2 and 3 (SI-5) are similar to those observed at strong and weak light intensities, respectively. We fitted the observed profiles using the sum of these contributions;

**Scheme 3.**
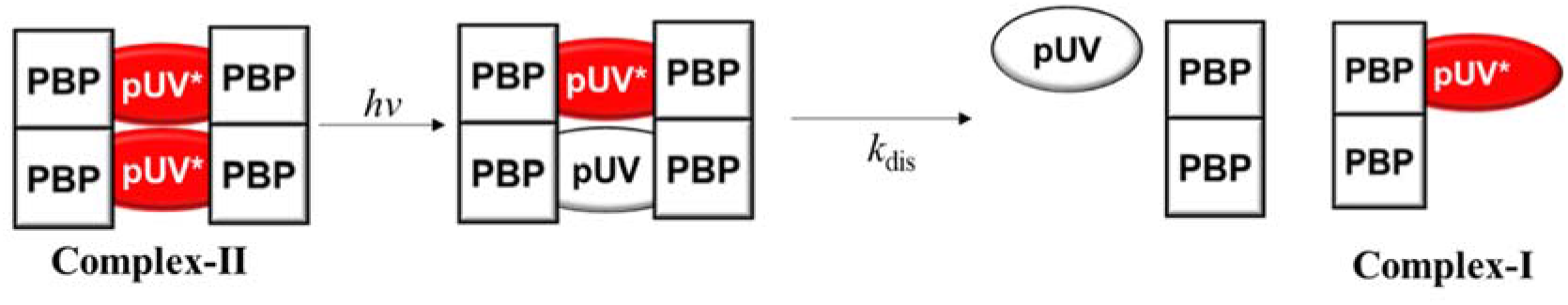

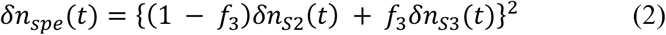

where f_3_ is the relative contribution from Scheme 3, and it is shown in Fig. 9(e) against the light intensity. A smaller f_3_ value upon increasing the light intensity is reasonable.

The reaction scheme determined in this study is illustrated in Fig. 10.

**Fig. 10.**
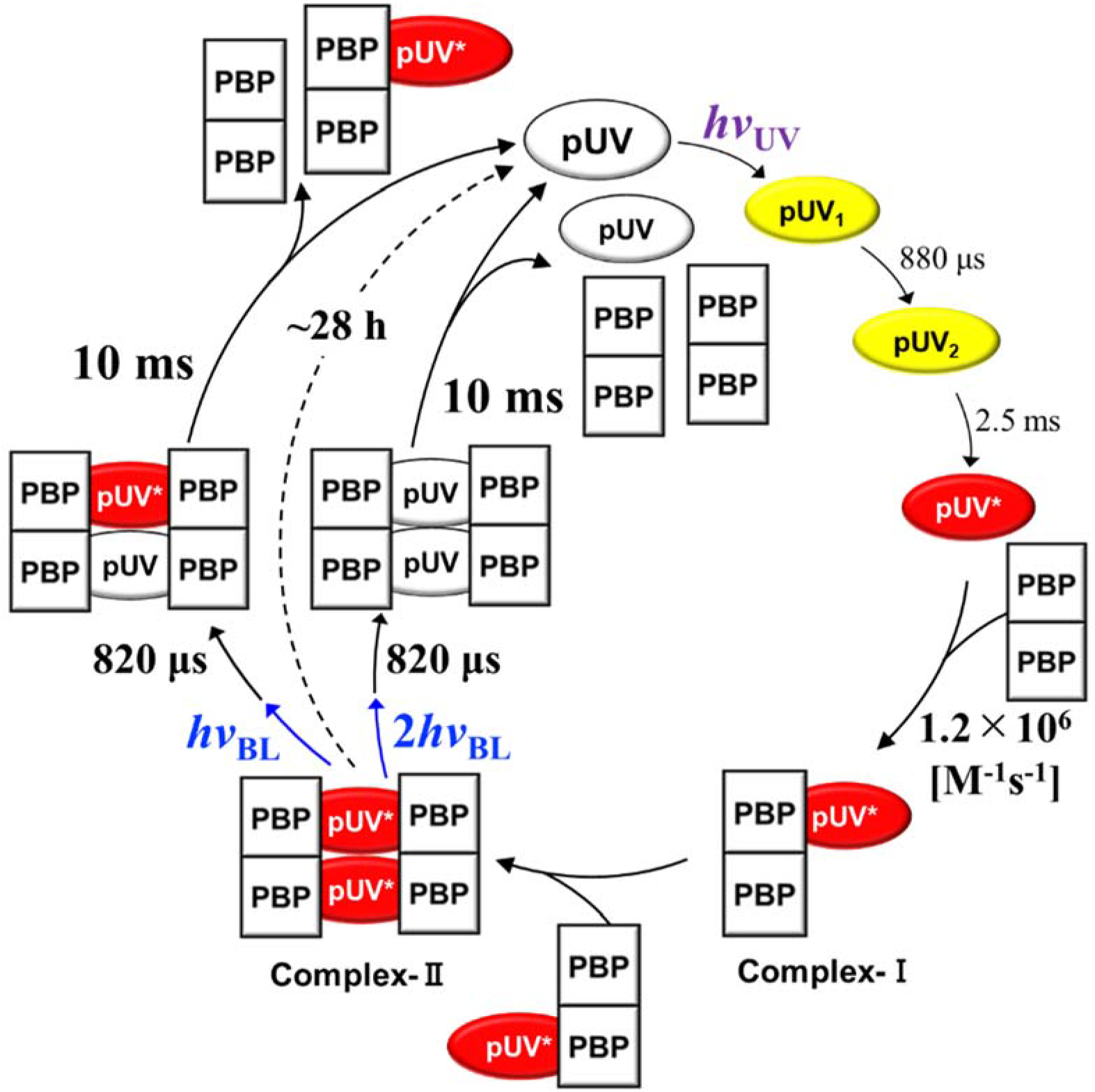
Schematic illustration of the reaction scheme for Rc-PYP with PBP. The intermediates before interacting with PBP are colored yellow and pUV* interacting with PBP is colored red.

## Discussion

It has been reported that the photoexcitation of Hh-PYP leads to the formation of pR and pB intermediates. It has been considered that pB, in which the chromophore is protonated and the N-terminal helices are unfolded, is considered to be the signaling state. In the case of Rc-PYP, similar characteristic features are observed for the reaction of pBL, that is, the chromophore is protonated and a conformation change is observed as the *D* changes in pBL_2_. This similarity suggests that pBL_2_ corresponds to pB in Hh-PYP and the possibility that pBL_2_ is the signaling state. In contrast to this consideration, in this study, we show that PBP does not interact with the intermediates in the pBL reaction. Hence, we believe that this blue light-absorbing species (pBL) is not responsible for signaling.

We demonstrated that the stable pUV* product interacts with PBP. Previously, it has been shown that the conformation change in one of the intermediates (pUV_2_) is larger from the viewpoint of the *D* value. If this intermediate interacts with PBP, this interaction is apparent as the diffusion signal changes in a fast time range. However, the *q*^2^-dependence of the diffusion signal is well reproduced by Scheme 1 (Fig. 4(b)). Hence, we consider that pUV*, not other short-lived intermediates, is the signaling state of Rc-PYP. A significant difference is the very long lifetime (~28 h) compared with that of pB for Hh-PYP (~1 s). In Hh-PYP, it is known that the lifetime of pB is controlled by the amino acid residues near the chromophore. In particular, mutations at M100 and Y98 drastically lengthen the lifetime of pB, for example, the lifetime of the M100A mutant is about 300 s. The long lifetime of pUV* may be due to the lack of M100, which is replaced by Gly in Rc-PYP. The long lifetime of pUV* and the similar long lifetime and stability of Complex-II should be beneficial for efficient signaling, which is important for biological function to avoid long-time exposure to harmful UV light. It may be interesting to note that Complex-II is formed from two Complex-I molecules, each of which is produced upon the photoexcitation of pUV, i.e. two photons are required to activate the UV response. Therefore, Rc-PYP can act as a non-linear light intensity sensor protein similar to previously reported proteins such as PixD and BlrP1.^27,28^ However, since the dark recovery of Complex-II is very slow, most of the pUV molecules may be excited even by weak UV light and the role of the non-linear light intensity dependence may not be important for biological function.

Previously, it has been shown that the relative fraction of pBL and pUV depends on the pH of the solution, i.e. the relative contribution from pUV increases upon decreasing the pH. Hence, the UV light sensitivity of Rc-PYP should depend on the pH. This observation suggests that Rc-PYP can act as a pH sensor using UV light.

One of the surprising features of the PYP-PBP interaction is that blue light completely recovers pUV* to the dark state in the presence of PBP, whereas slight pUV* formation is detected upon BL(480) irradiation in the absence of PBP. Previously, this minor change in the absence of PBP was explained in terms of a possible side reaction from the excited state of pBL or a stepwise multi-excitation from excited pBL.^15^ The different behavior observed in the presence and absence of PBP suggests that the efficiency of the blue light induced dark recovery process increases in the presence of PBP.

The dissociation reaction of Complex-II is mainly studied by the diffusion detection method. The excitation light intensity dependence of the diffusion signal suggests that the excitation of two pUV* molecules leads to complete dissociation and the excitation of one pUV* molecule produces pUV, PBP, and Complex-I. However, it is possible that the excitation of one pUV* molecule leads to the complete dissociation of pUV, pUV*, and two PBP dimers. Even in this case, Complex-I can be produced from pUV* and PBP with a time constant of ~8 ms, which is faster than the time range of the diffusion signal. Hence, we cannot exclude the possibility of complete dissociation. However, we considered that the complete dissociation scheme upon the excitation of one pUV* molecule is possible, but less plausible, because Complex-I is stable before forming Complex-II. In any case, Complex-I should reproduce Complex-II upon binding with another Complex-I, which is created during the dissociation reaction. The rate of this process should be concentration dependent and may be too slow to be detected in this study. Hence, this process is ignored in the reaction scheme shown in Fig. 10.

The isoelectric points of Rc-PYP and PBP are calculated to be 9.4 and 3.9 from their amino acid sequences. This large difference suggests that the electrostatic binding could be important in their intermolecular interactions. However, Rc-PYP in the dark state does not form a stable complex with PBP. It is likely that the conformation change in pUV* is necessary to expose the electrostatic binding site for its association with PBP.

One significant finding in this study is that Rc-PYP interacts with PBP upon irradiation with UV light. Therefore, we propose that Rc-PYP is a UV sensor protein. Not many UV sensor proteins have been known so far. A typical example is UV Resistance Locus 8 (UVR8), which was initially identified from *Arabidopsis thaliana* as a photoreceptor that responds to ultraviolet-B (UV-B) light.^29^ UVR8 is a relatively large protein (~40 kDa) without any chromophore besides amino acid residues bearing aromatic side chains, and forms a dimer in its native state.^29^ The dimer should be dissociated into the monomers for interacting with the downstream partner protein, COP1.^30^ Compared with the rather complex structure of UVR8,^31^ Rc-PYP is a simple and monomeric protein. Its interaction with the downstream protein is controlled by conformation changes. In this respect, Rc-PYP is a new type of UV sensor, of which signaling process is similar to that of other light sensor proteins in the visible light region.

It may be interesting to note that, even for well-studied Hh-PYP, an absorption band in the UV region appears at low pH.^32^ The spectrum of this UV band is similar to that of pUV. This similarity suggests a possibility that PBP can interact with Hh-PYP upon UV excitation under low pH conditions. We examined this possibility by the diffusion detection method. We found that Hh-PYP does not interact with PBP under low or neutral pH conditions. Furthermore, a protein with a similar sequence to PBP is not found in the *Halorhodospira halophila* genome. Therefore, the downstream protein for Hh-PYP should have a sequence different to that of PBP for Rc-PYP.

Finally, it should be mentioned that this Rc-PYP and PBP system can be used as a novel optogenetic tool. The very long lifetime of Complex-II results in high sensitivity and the switchable reaction depending on the wavelength of light will be useful toward controlling the reactions.

## Conclusions

In this study, the intermolecular interaction dynamics between photoexcited Rc-PYP and a possible downstream target protein, PBP are investigated in the time domain. It was found that pBL and any intermediates formed during the pBL reaction do not interact with PBP. Only the long-lived pUV* product, is found to interact with PBP. We determined the rates of the association and dissociation reactions in a series of interaction schemes. Using the time-resolved diffusion method, we found that pUV* first interacts with one PBP dimer to produce Complex-I at a rate of 1.2 × 10^6^ M^−1^s^−1^ upon the photoexcitation of pUV. Subsequently, two Complex-I molecules associate to form Complex-II, which is a long-lived species and this process is spectrally silent. The size of the complexes depends on the relative concentrations of PYP and PBP, and the light intensity. In the dissociation process, pUV* in Complex-II first recovers to pUV with a time constant of 820 μs and the dissociation to form pUV and PBP dimer takes place within 10 ms. It should be noted that UV light induces the intermolecular interaction between Rc-PYP and PBP, and the blue light switches off this interaction. Therefore, Rc-PYP is a unique photochromic UV light sensor.

## ACKNOWLEDGEMENTS

This work was supported by a Grant-in-aid for Scientific Research on Innovative Areas (research in a proposed research area) (Nos. JP20107003, and JP25102004 to M.T.) and a Grant-in-aid for Scientific Research from MEXT/JSPS (25288005, 17H03008, 19H01863 to M.T., 17H05001 and 20H04708 to Y.N.), and the Sasakawa Scientific Research Grant (2020-3020 to S.K.).

## References

(1) Sprenger, W. W.; Hoff, W. D.; Armitage, J. P.; Hellingwerf, K. J. The Eubacterium Ectothiorhodospira Halophila Is Negatively Phototactic, with a Wavelength Dependence That Fits the Absorption Spectrum of the Photoactive Yellow Protein. J. Bacteriol. 1993, 175 (10), 3096–3104.

(2) Imamoto, Y.; Kataoka, M. Structure and Photoreaction of Photoactive Yellow Protein, a Structural Prototype of the PAS Domain Superfamily†. Photochem. Photobiol. 2007, 83 (1), 40–49.

(3) Ramachandran, P. L.; Lovett, J. E.; Carl, P. J.; Cammarata, M.; Lee, J. H.; Jung, Y. O.; Ihee, H.; Timmel, C. R.; van Thor, J. J. The Short-Lived Signaling State of the Photoactive Yellow Protein Photoreceptor Revealed by Combined Structural Probes. J. Am. Chem. Soc. 2011, 133 (24), 9395–9404.

(4) Mix, L. T. Diversity in the Photodynamics of the Photoactive Yellow Protein Family; University of California, Davis, 2018.

(5) Kuramochi, H.; Takeuchi, S.; Yonezawa, K.; Kamikubo, H.; Kataoka, M.; Tahara, T. Probing the Early Stages of Photoreception in Photoactive Yellow Protein with Ultrafast Time-Domain Raman Spectroscopy. Nat. Chem. 2017, 9 (7), 660–666.

(6) Pande, K.; Hutchison, C. D. M.; Groenhof, G.; Aquila, A.; Robinson, J. S.; Tenboer, J.; Basu, S.; Boutet, S.; DePonte, D. P.; Liang, M.; White, T. A.; Zatsepin, N. A.; Yefanov, O.; Morozov, D.; Oberthuer, D.; Gati, C.; Subramanian, G.; James, D.; Zhao, Y.; Koralek, J.; Brayshaw, J.; Kupitz, C.; Conrad, C.; Roy-Chowdhury, S.; Coe, J. D.; Metz, M.; Xavier, P. L.; Grant, T. D.; Koglin, J. E.; Ketawala, G.; Fromme, R.; Šrajer, V.; Henning, R.; Spence, J. C. H.; Ourmazd, A.; Schwander, P.; Weierstall, U.; Frank, M.; Fromme, P.; Barty, A.; Chapman, H. N.; Moffat, K.; van Thor, J. J.; Schmidt, M. Femtosecond Structural Dynamics Drives the Trans/Cis Isomerization in Photoactive Yellow Protein. Science 2016, 352 (6286), 725–729.

(7) Meyer, T. E.; Yakali, E.; Cusanovich, M. A.; Tollin, G. Properties of a Water-Soluble, Yellow Protein Isolated from a Halophilic Phototrophic Bacterium That Has Photochemical Activity Analogous to Sensory Rhodopsin. Biochemistry 1987, 26 (2), 418–423.

(8) Hoff, W. D.; van Stokkum, I. H.; van Ramesdonk, H. J.; van Brederode, M. E.; Brouwer, A. M.; Fitch, J. C.; Meyer, T. E.; van Grondelle, R.; Hellingwerf, K. J. Measurement and Global Analysis of the Absorbance Changes in the Photocycle of the Photoactive Yellow Protein from Ectothiorhodospira Halophila. Biophys. J. 1994, 67 (4), 1691–1705.

(9) Kay, S. A. PAS, Present, and Future: Clues to the Origins of Circadian Clocks. Science 1997, 276(5313), 753–754.

(10) Huang, Z. J.; Edery, I.; Rosbash, M. PAS Is a Dimerization Domain Common to Drosophila Period and Several Transcription Factors. Nature 1993, 364 (6434), 259–262.

(11) McGuire, J.; Coumailleau, P.; Whitelaw, M. L.; Gustafsson, J. A.; Poellinger, L. The Basic Helix-Loop-Helix/PAS Factor Sim Is Associated with Hsp90. Implications for Regulation by Interaction with Partner Factors. J. Biol. Chem. 1995, 270 (52), 31353–31357.

(12) Pellequer, J. L.; Wager-Smith, K. A.; Kay, S. A.; Getzoff, E. D. Photoactive Yellow Protein: A Structural Prototype for the Three-Dimensional Fold of the PAS Domain Superfamily. Proc. Natl. Acad. Sci. U. S. A. 1998, 95 (11), 5884–5890.

(13) Khan, J. S.; Imamoto, Y.; Yamazaki, Y.; Kataoka, M.; Tokunaga, F.; Terazima, M. A Biosensor in the Time Domain Based on the Diffusion Coefficient Measurement: Intermolecular Interaction of an Intermediate of Photoactive Yellow Protein. Anal. Chem. 2005, 77 (20), 6625–6629.

(14) Kyndt, J. A.; Hurley, J. K.; Devreese, B.; Meyer, T. E.; Cusanovich, M. A.; Tollin, G.; Van Beeumen, J. J. Rhodobacter Capsulatus Photoactive Yellow Protein: Genetic Context, Spectral and Kinetics Characterization, and Mutagenesis. Biochemistry 2004, 43 (7), 1809–1820.

(15) Kim, S.; Nakasone, Y.; Takakado, A.; Yamazaki, Y.; Kamikubo, H.; Terazima, M. Wavelength-Dependent Photoreaction of PYP from Rhodobacter Capsulatus. Biochemistry 2020, 59 (51), 4810–4821.

(16) Strnad, H.; Lapidus, A.; Paces, J.; Ulbrich, P.; Vlcek, C.; Paces, V.; Haselkorn, R. Complete Genome Sequence of the Photosynthetic Purple Nonsulfur Bacterium Rhodobacter Capsulatus SB 1003. J. Bacteriol. 2010, 192 (13), 3545–3546.

(17) Drozdetskiy, A.; Cole, C.; Procter, J.; Barton, G. J. JPred4: A Protein Secondary Structure Prediction Server. Nucleic Acids Res. 2015, 43 (W1), W389–94.

(18) Imamoto, Y.; Koshimizu, H.; Mihara, K.; Hisatomi, O.; Mizukami, T.; Tsujimoto, K.; Kataoka, M.; Tokunaga, F. Roles of Amino Acid Residues near the Chromophore of Photoactive Yellow Protein. Biochemistry 2001, 40 (15), 4679–4685.

(19) Gasteiger, E.; Hoogland, C.; Gattiker, A.; Duvaud, S.; Wilkins, M. R.; Appel, R. D.; Bairoch, A. Protein Identification and Analysis Tools on the ExPASy Server. In The Proteomics Protocols Handbook; Walker, J. M., Ed.; Humana Press: Totowa, NJ, 2005; pp 571–607.

(20) Terazima, M. Diffusion Coefficients as a Monitor of Reaction Kinetics of Biological Molecules. Phys. Chem. Chem. Phys. 2006, 8 (5), 545–557.

(21) Terazima, M. Spectrally Silent Protein Reaction Dynamics Revealed by Time-Resolved Thermodynamics and Diffusion Techniques. Acc. Chem. Res. 2021, 54 (9), 2238–2248.

(22) Terazima, M. Time-Dependent Intermolecular Interaction during Protein Reactions. Phys. Chem. Chem. Phys. 2011, 13 (38), 16928–16940.

(23) Terazima, M. Studies of Photo-Induced Protein Reactions by Spectrally Silent Reaction Dynamics Detection Methods: Applications to the Photoreaction of the LOV2 Domain of Phototropin from Arabidopsis Thaliana. Biochimica et Biophysica Acta (BBA) - Proteins and Proteomics 2011, 1814 (8), 1093–1105.

(24) Nakasone, Y.; Zikihara, K.; Tokutomi, S.; Terazima, M. Kinetics of Conformational Changes of the FKF1-LOV Domain upon Photoexcitation. Biophys. J. 2010, 99 (11), 3831–3839.

(25) Nakasone, Y.; Kikukawa, K.; Masuda, S.; Terazima, M. Time-Resolved Study of Interprotein Signaling Process of a Blue Light Sensor PapB–PapA Complex. J. Phys. Chem. B 2019, 123 (15), 3210–3218.

(26) Takeda, K.; Terazima, M. Dynamics of Conformational Changes in Full-Length Phytochrome from Cyanobacterium Synechocystis Sp. PCC6803 (Cph1) Monitored by Time-Resolved Translational Diffusion Detection. Biochemistry 2019, 58 (24), 2720–2729.

(27) Tanaka, K.; Nakasone, Y.; Okajima, K.; Ikeuchi, M.; Tokutomi, S.; Terazima, M. Light-Induced Conformational Change and Transient Dissociation Reaction of the BLUF Photoreceptor Synechocystis PixD (Slr1694). J. Mol. Biol. 2011, 409 (5), 773–785.

(28) Shibata, K.; Nakasone, Y.; Terazima, M. Photoreaction of BlrP1: The Role of a Nonlinear Photo-Intensity Sensor. Phys. Chem. Chem. Phys. 2018, 20 (12), 8133–8142.

(29) Rizzini, L.; Favory, J.-J.; Cloix, C.; Faggionato, D.; O’Hara, A.; Kaiserli, E.; Baumeister, R.; Schäfer, E.; Nagy, F.; Jenkins, G. I.; Ulm, R. Perception of UV-B by the Arabidopsis UVR8 Protein. Science 2011, 332 (6025), 103–106.

(30) Miyamori, T.; Nakasone, Y.; Hitomi, K.; Christie, J. M.; Getzoff, E. D.; Terazima, M. Reaction Dynamics of the UV-B Photosensor UVR8. Photochem. Photobiol. Sci. 2015, 14 (5), 995–1004.

(31) Christie, J. M.; Arvai, A. S.; Baxter, K. J.; Heilmann, M.; Pratt, A. J.; O’Hara, A.; Kelly, S. M.; Hothorn, M.; Smith, B. O.; Hitomi, K.; Jenkins, G. I.; Getzoff, E. D. Plant UVR8 Photoreceptor Senses UV-B by Tryptophan-Mediated Disruption of Cross-Dimer Salt Bridges. Science 2012, 335(6075), 1492–1496.

(32) Meyer, T. E. Isolation and Characterization of Soluble Cytochromes, Ferredoxins and Other Chromophoric Proteins from the Halophilic Phototrophic Bacterium Ectothiorhodospira Halophila. Biochimica et Biophysica Acta (BBA) - Bioenergetics 1985, 806 (1), 175–183.

