## Supporting Information for "Interaction of a photochromic UV sensor protein Rc-PYP with PYP-binding protein"

### SI-1. Amino acid sequence and secondary structure of PYP-Binding protein (PBP)

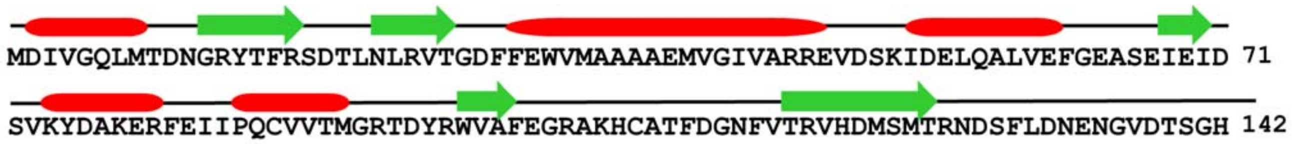


Fig. S1 The amino acid residues and predicted secondary structural of PBP by the Jpred4 software. The α-helices and β-sheets are shown as red and green, respectively.

### SI-2. Dark recovery after UV irradiation

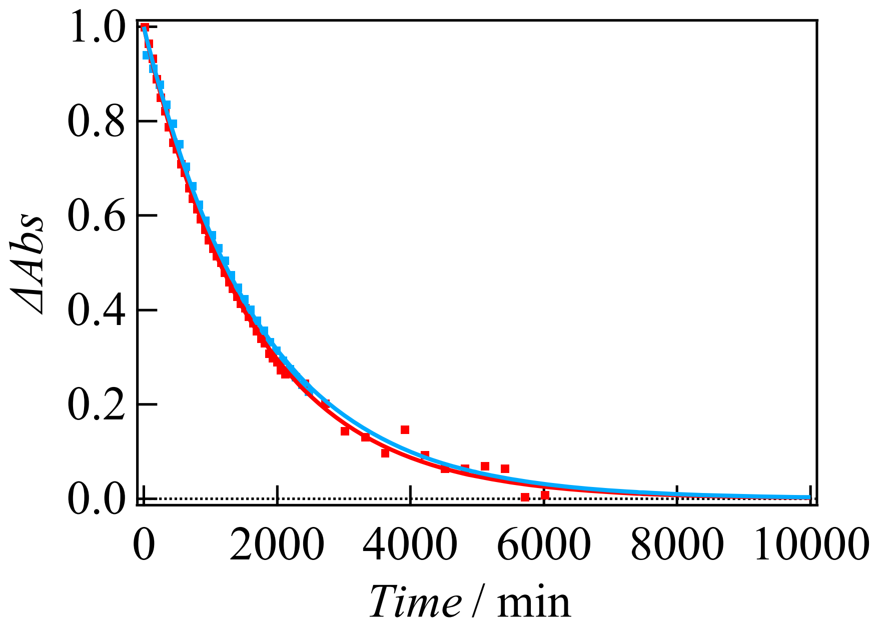

Fig. S2. The dark recovery kinetics of Rc-PYP at 10 μM without (red dots) and with PBP at 100 μM (blue dots) monitored by the absorption at 438 nm. Absorption intensities are normalized by the absorption differences between the dark and light states. The best fitted curves by a single-exponential function (smooth curves) are also shown.

**SI-3. Analytical equation of the diffusion signal based on Scheme1**

The time profile of the species grating component based on scheme 1 ($\delta n_{S1}(t)$) may be expressed as;

$$\begin{aligned}
\delta n_{S1}(t) = & -\delta n_{UV}\exp(-D_{UV}q^2t) + \delta n_{UV2}\exp\{-(D_{UV2}q^2+k_2)t\} \\
& + \frac{k_2\delta n_{UV*}}{(D_{UV2}-D_{UV*})q^2+k_2-k_3}[\exp\{-(D_{UV*}q^2+k_3)t\} - \exp\{-(D_{UV2}q^2+k_2)t\}] \\
& + \frac{k_2k_3\delta n_{C1}}{(D_{UV2}-D_{UV*})q^2+k_2-k_3}[\frac{1}{(D_{UV2}-D_{c1})q^2+k_2}exp\{-(D_{UV2}q^2+k_2)t\} \\
& - \frac{1}{(D_{UV*}-D_{c1})q^2+k_3}exp\{-(D_{UV*}q^2+k_3)t\} \\
& + \left\{\frac{1}{(D_{UV*}-D_{c1})q^2+k_3} - \frac{1}{(D_{UV2}-D_{c1})q^2+k_2}\right\}exp\{-D_{C1}q^2t\}] \\
& - \delta n_{PBP}\frac{k_2k_3}{(D_{UV2}-D_{UV*})q^2+k_2-k_3}[-\frac{1}{(D_{UV2}-D_{PBP})q^2+k_2}\{exp(-D_{PBP}q^2t) \\
& - exp\{-(D_{UV2}q^2+k_2)t\}\} \\
& + \frac{1}{(D_{UV*}-D_{PBP})q^2+k_3}\{exp(-D_{PBP}q^2t) - exp\{-(D_{UV*}q^2+k_3)t\}\}]
\end{aligned}$$

(S1)

where $\delta n_i$ and $D_i$ (i=UV, UV2, UV*, C1, and PBP) denote the refractive index changes and diffusion coefficients of the i-species, respectively. The subscripts of UV, UV2, UV*, C1, and PBP represent the species of pUV, $pUV_2$, pUV*, Complex-I, $PBP_2$, respectively.

**SI-4. Analytical equation of the diffusion signal based on Scheme 2 and Scheme 3.**

The time profile of the species grating component based on scheme 2 ($\delta n_{S2}(t)$) is given as;

$$\begin{aligned}
\delta n_{S2}(t) = & -\delta n_{C2}\exp(-D_{C2}q^2t) + \delta n_{C2'}\exp\{-(D_{C2'}q^2+k_{dis})\mathrm{t}\} \\
& + \frac{2\delta n_{UV}k_{dis}}{(D_{UV}-D_{C2'})q^2-k_{dis}}[\exp\{-(D_{C2'}q^2+k_{dis})\mathrm{t}\} - \exp(-D_{UV}q^2t)] \\
& + \frac{2\delta n_{PBP}k_{dis}}{(D_{PBP}-D_{C2'})q^2-k_{dis}}[\exp\{-(D_{C2'}q^2+k_{dis})\mathrm{t}\} - \exp(-D_{PBP}q^2t)]
\end{aligned}$$

(S2)

where $\delta n_i$ and $D_i$ (i=C2, C2', UV, and PBP) denote the refractive index changes and diffusion coefficients of the i-species, respectively. The subscripts of C2 and C2' represent the species of Complex-II, the complex of pUV$_2$-PBP$_4$, respectively.

The time profile of the species grating component based on scheme 3 ($\delta n_{S3}(t)$) is given as;

$$\delta n_{S3}(t) = -\delta n_{C2} \exp(-D_{C2} q^2 t) \mp \delta n_{C2''} \exp\{-(D_{C2''} q^2 + k_{dis})t\}$$
$$+ \frac{\delta n_{UV} k_{dis}}{(D_{UV} - D_{C2''})q^2 - k_{dis}} [\exp\{-(D_{C2''} q^2 + k_{dis})t\} - \exp(-D_{UV} q^2 t)]$$
$$+ \frac{\delta n_{PBP} k_{dis}}{(D_{PBP} - D_{C2''})q^2 - k_{dis}} [\exp\{-(D_{C2''} q^2 + k_{dis})t\} - \exp(-D_{PBP} q^2 t)]$$
$$+ \frac{\delta n_{c1} k_{dis}}{(D_{c1} - D_{C2''})q^2 - k_{dis}} [\exp\{-(D_{C2''} q^2 + k_{dis})t\} - \exp(-D_{c1} q^2 t)] \qquad \text{(S3)}$$

where $\delta n_{C2''}$ and $D_{C2''}$ denote the refractive index changes and diffusion coefficients of the complex of pUV-pUV*-PBP$_4$ in Scheme 3.

**SI-5. Simulation of TG signal in dissociation process.**

The TG signals based on Scheme 2 and Scheme 3 are calculated by eq.(S2) and (S3). The parameters in eq.(S2) was fixed to the value determined by the analysis of $q^2$ dependence in main text. For calculation of eq.(S3), the same values as the calculation of eq.(S2), $\delta n_{C2}=\delta n_{C2''}$, and $D_{C2}=D_{C2''}$ are used.

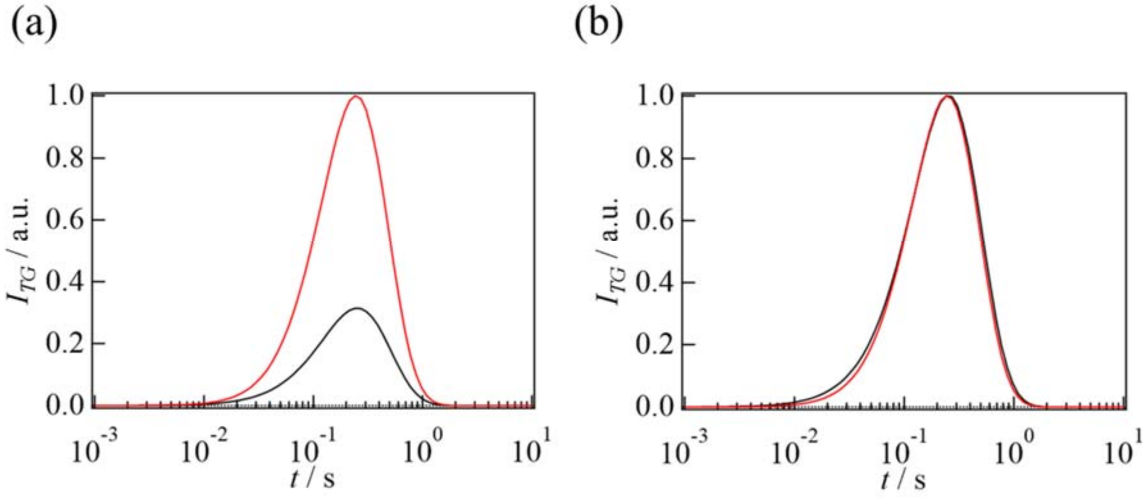


Figure S3. (a) Simulated TG signals based on eq.(S2) (red line) and eq.(S3) (black line). (b) The signals normalized at the peak intensities.
